# Modeling the 3D structure and conformational dynamics of very large RNAs using coarse-grained molecular simulations

**DOI:** 10.1101/2023.06.06.543892

**Authors:** Aaron N. Henderson, Robert T. McDonnell, Adrian H. Elcock

**Affiliations:** Department of Biochemistry & Molecular Biology, University of Iowa, Iowa City, Iowa, USA

## Abstract

We describe a computational approach to building and simulating realistic 3D models of very large RNA molecules (>1000 nucleotides) at a resolution of one “bead” per nucleotide. The method starts with a predicted secondary structure and uses several stages of energy minimization and Brownian dynamics (BD) simulation to build 3D models. A key step in the protocol is the temporary addition of a 4^th^ spatial dimension that allows all predicted helical elements to become disentangled from each other in an effectively automated way. We then use the resulting 3D models as input to Brownian dynamics simulations that include hydrodynamic interactions (HIs) that allow the diffusive properties of the RNA to be modelled as well as enabling its conformational dynamics to be simulated. To validate the dynamics part of the method, we first show that when applied to small RNAs with known 3D structures the BD-HI simulation models accurately reproduce their experimental hydrodynamic radii (Rh). We then apply the modelling and simulation protocol to a variety of RNAs for which experimental Rh values have been reported ranging in size from 85 to 3569 nucleotides. We show that the 3D models, when used in BD-HI simulations, produce hydrodynamic radii that are usually in good agreement with experimental estimates for RNAs that do not contain tertiary contacts that persist even under very low salt conditions. Finally, we show that sampling of the conformational dynamics of large RNAs on timescales of 100 µs is computationally feasible with BD-HI simulations.

## Introduction

The central importance of RNA to many biological processes has meant that many computational methods have been developed with the aim of predicting the 3D structures of RNA molecules. Most of the available methods focus on RNAs that contain at most ∼250 nucleotides (nts) and that attempt to predict their structures with atomic levels of detail (see, e.g. (1–9)). Very recently, a number of deep learning-based structure prediction methods have been reported (10–13), and it is likely that with the proven success of AlphaFold2 in the realm of protein structure prediction (14,15), further developments for RNAs of this size will follow in the near future. In a biological setting, however, many RNAs far exceed 1000 nts in size: in *E. coli*, for example, mRNAs can contain as many as 17,000 nts, while in higher organisms long non-coding RNAs (lncRNAs) can have lengths in excess of 200,000 nts (16). When large RNAs adopt stable 3D structures, conventional experimental techniques can be used to obtain high resolution structural models. For *E. coli*, for example, high resolution structures of ribosomes have been available for many years (e.g. (17–19)), and more recently, cryoEM techniques have been used to obtain structures of ribosomal subunit assembly intermediates (e.g. (20,21)). For viral genomes, there is also the possibility of solving structures *in situ*: medium-resolution and partial atomic structures of the encapsidated MS2 genome, for example, were reported a few years ago (22,23) and a complete atomic model of the encapsidated 4127-nt Qβ phage genome has been reported recently (24); efforts such as these can be facilitated by automated methods for modeling RNA structures into cryoEM data (e.g. (25,26)). When large RNAs do not adopt stable folded forms, lower-resolution experimental methods such as small-angle X-ray scattering (SAXS) can be used, often in combination with modelling methods, to produce plausible but more coarse-grained models of large RNAs (27,28); SAXS measurements have even been used to visualize the encapsidated MS2 genome (29). Alternatively, insights into secondary structure branching patterns – and the wide range of different secondary structures that can be simultaneously populated – can be obtained by direct cryoEM imaging of large RNAs in solution (28,30).

While a number of very large RNA models have been produced over the years – e.g. the Harvey group’s groundbreaking atomic model of the entire STMV virus with its 1058-nt RNA genome (31), Poblete & Guzman’s interesting comparative CG models of the same genome in its *in virio* and *in vitro* forms (32) and, noted above, the Zhang group’s report of a complete atomic model of the encapsidated Qβ genome (24) – very few of the methods developed to predict RNA structure have been consistently used to model RNAs that are much larger than 250 nts. Two of these methods, RNAComposer (33) and 3dRNA (6) are accessible as webservers and produce atomically detailed models, while the third, NAST (34), is a stand-alone program that produces coarse-grained (CG) models with a resolution of one bead per nucleotide. In common with many other RNA structure prediction methods, all of these methods start with an assumed secondary structure as input. Both RNAComposer and 3dRNA produce models by piecing together structural fragments extracted from solved RNA structures; the resulting models are often highly plausible, and importantly, are returned rapidly, i.e. within a few minutes in the case of RNAComposer. However, our own experience with applying both methods to larger RNAs is that the predicted models can sometimes contain instances of unphysically “entangled” helices, i.e. cases in which the backbone of one strand of a double-helical segment passes directly through the middle of a second double-helical segment. A recent study by the Szachniuk group has shown that such entanglements occur with quite high frequency (∼13 % of the time) in RNA structure prediction scenarios, thereby indicating that this problem is not unique to RNAComposer and 3dRNA (35).

In contrast to the above two methods, NAST (Nucleic Acid Simulation Tool) (34), developed by the Altman group employs a coarse-grained (CG) force field designed to be used in combination with molecular dynamics simulations to produce RNA models. Starting from an initial model that is either completely linear or circular, force field terms are applied to bring base-paired nucleotides together to form double helical regions specified in the predicted secondary structure. While it was only tested on RNAs up to 158 nts in the original publication, subsequent work from the Gelbart group used NAST to produce 3D models of RNAs up to 2777 nts in length (28). One significant drawback that we have found in our own attempts to use NAST is its choice of initial model: neither a linear nor a circular model are particularly good starting points for RNAs that contain large numbers of double-stranded regions, and formation of the base-pairing interactions (which requires widely-separated nucleotides to travel very long distances) can result in significant conformational strain in the final models.

To address each of the issues outlined above, we report here the development of a computational protocol suitable for building and simulating models of large RNAs, which we define here as RNAs containing ∼1000 nts or more. Throughout, we use a simulation-based approach to build models of RNAs, so the philosophy used here is very similar to that underpinning NAST (34). There are, however, a number of crucial differences. First, we attempt to establish a better initial model for the process: instead of starting from a linear or circular model, we start with a model that has all the predicted secondary structure elements pre-formed. Second, to remove the helical entanglements that typically result when very large RNA models are built (see above), we implement a novel simulation approach that we developed recently (36); this approach makes use of a transiently added 4^th^ spatial dimension that allows secondary structure elements to be moved apart from each other automatically. Third, having built complete 3D models, we use Brownian dynamics (BD) simulations that include hydrodynamic interactions (HIs) to model the RNA’s diffusive and conformational dynamics; simulations of this type have been shown by us previously to perform well for proteins (37) and, by the Garcia de la Torre group, for structured RNAs (38). Importantly, the ability to realistically model the diffusive behavior of large RNAs permits us to make direct comparison with experimental hydrodynamic radius data (Rh) and to obtain molecular level views of the timescales of internal conformational dynamics. The remainder of this manuscript describes in detail the method, which we call *uiowa_rna*, before outlining its application to a variety of RNA systems ranging from small 85 nt RNAs, to RNAs containing >3500 nts.

## Materials & Methods

### The RNA Model

Comprehensive reviews of the many possible ways to develop and apply CG models of RNAs have been reported recently by the Bujnicki (39) and Chen groups (40). *uiowa_rna* seeks to build CG models of RNAs in which each CG bead represents a single nucleotide; this level of resolution has been chosen to match that adopted in our previous model of the complete *E. coli* chromosome (41), and is the same as that used, for example, in a BD-HI simulation study of two RNAs reported by the Garcia de la Torre group (38), in the pioneering efforts to model large RNAs conducted by the Harvey group (42), and to model unfolding and refolding pathways in RNAs by the Thirumalai group (43). Here, the representative bead for each nucleotide is placed at the position occupied by the phosphorous atom. Our final 3D models are constructed following a stepwise protocol that is outlined in Figure 1.

**Figure 1.**
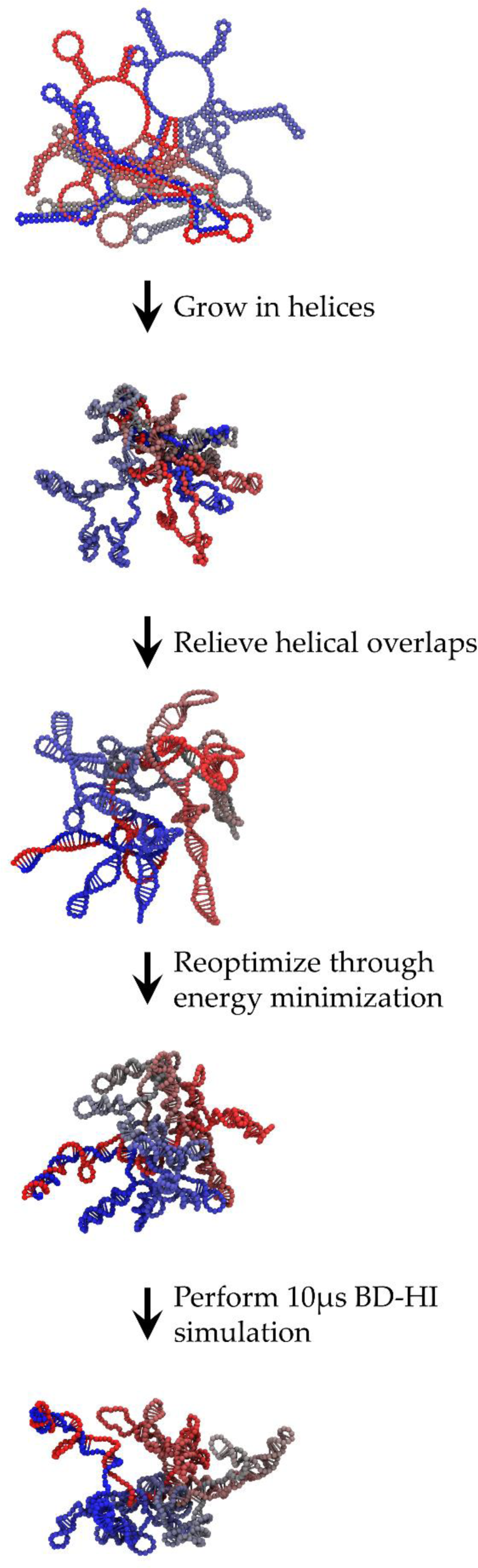
Illustration of the 3D modeling protocol. Images show the progressive inclusion of 3D structure to a model of the 961-nt BunVS RNA starting from its predicted secondary structure to its structure at the end of a 10 µs BD-HI simulation. At each stage of the protocol, the RNA is colored with a blue-gray-red gradient starting at the 5’ end. These and all other images of molecular models in this manuscript were made with VMD (86).

### Secondary Structure Prediction

The starting point for the protocol presented here is the RNA’s secondary structure. While this can increasingly be determined via experimental measurements (e.g. (44–47)) it probably remains more usual for it to be predicted computationally. In this work, two methods of secondary structure prediction have been used: RNAFold (48) and CoFold (49). RNAFold is part of the widely used ViennaRNA package (version 2.4.1), and as typically used here, seeks to find the minimum free energy (MFE) secondary structure using the energy parameters developed and assembly by the Turner and Mathews groups (50,51). CoFold is an extension of an earlier iteration of RNAFold (that found in ViennaRNA package 2.0), that seeks to penalize long-range base-pairing interactions relative to short-range interactions in an attempt to more accurately predict the potential effects of cotranscriptional folding. Because of this, the secondary structures predicted using the two methods for very large RNAs can be very different (see Results). Most of the models reported here were based on predictions of the MFE secondary structure obtained with the two programs; to test the effects of using different secondary structures as input to our 3D modelling protocol, however, additional models of two selected large RNAs were built using thermally accessible secondary structures randomly sampled by RNAFold using the command-line option “--ImFeelingLucky” (see Results). In all other cases, RNAFold and CoFold were run using their default parameters with the only exception being that we explicitly disallowed the formation of double-helical segments that contained only a single base-pair (using the option “--noLP”).

For the specific case of the 16S rRNA, we also made use of an additional secondary structure prediction algorithm provided by the Vienna Package (52). This is the program RNAcofold (53) (not to be confused with CoFold), which is designed to predict the secondary structure of dimeric RNAs formed from the non-covalent association of two RNA monomers. The MFE structure of a 16S rRNA dimer was predicted using RNAcofold’s default parameters.

### Tertiary Structure Prediction

With a secondary structure generated, the *uiowa_rna* protocol begins by constructing an initial idealized 3D model. This is achieved by first using the ViennaRNA utility RNAplot (50-52,54,55) to make a 2D postscript image of the secondary structure (using the “simple radial layout” option invoked using the flag “-t0”) and then converting it into a .pdb format file with a script written in-house. The resulting structure (see top image in Figure 1) serves as a starting point for a series of subsequent BD simulations that seek to add in the 3D details. These simulations are conducted in four stages, with a generic molecular mechanics-based approach being used throughout the process to model the RNA’s conformational behavior: covalent bonds are placed between CG beads that are adjacent in sequence or that are base-paired to each other, angle terms are used to describe bending deformations of three consecutive bonded CG beads (e.g. i:i+1:i+2 beads), and dihedral terms are used to describe torsional degrees of freedom involving four consecutive bonded CG beads (e.g. i:i+1:i+2:i+3 beads). The force constants applied to these degrees of freedom, and the timesteps used in the BD simulations, vary somewhat with the stage of the protocol.

The first stage of the protocol involves “growing in” 3D structure into the initially flat, ladder-like double helical segments of the RNA. This is achieved by performing a series of short simulations with our BD simulation code, *uiowa_bd* (56), over the course of which all internal parameters (i.e. equilibrium bond lengths, angles, and dihedrals) are interpolated from their values in the initial model to the values expected in an idealized A-form RNA double helix. The internal parameters of the latter were obtained from a model of a C_40_:G_40_ double helix generated using the online make_NA-server (57). Throughout this stage of the process all bonds are restrained to their target values using a simple harmonic potential function with a force constant of 20 kcal/mol/Å. To give priority to the formation of correct geometries in double-helical regions over single-stranded regions stronger force constants are applied to those CG beads that are involved in base-pairing interactions: force constants for the angle terms are set to 5.0 and 2.5 kcal/mol/rad, for base-paired and single-stranded CG beads respectively, while the barrier heights of their dihedral energy terms are set to 1 and 0.1 kcal/mol, respectively. We use the same sets of equilibrium dihedral angles for single-stranded regions of the RNA in lieu of more complicated knowledge-based energy terms since structures reported by the Ono and Kim groups have indicated that a general double-helical nature is often retained even when there are non-base pairing nucleotides in the helix (58,59). Other than the bonded interactions, no van der Waals or electrostatic interactions are included at this stage of the protocol. Each of the BD simulations is carried out for 1 ns using a timestep of 50 fs. The second image in Figure 1 depicts a typical output structure from this simulation stage: the model now contains double helical segments that have the expected appearance of conventional A-RNA, but owing to the absence of steric interactions up to this point, many CG beads overlap with one another.

The second stage of the protocol seeks to simultaneously introduce steric interactions and to resolve potential helical entanglements. It does so using a novel simulation approach (36) in which a 4^th^ spatial dimension is temporarily introduced and then eventually compressed out of existence (see below). The addition of an extra spatial dimension to the simulations allows pairs of CG beads that would otherwise be considered clashing in 3D to be placed in a non-clashing arrangement by simply providing them with suitably spaced coordinates in the added dimension. The assignment of these initial displacements in the 4^th^ spatial dimension is achieved as follows. First, a simple computational analysis of the list of base-pairs is used to assign all nucleotides to either double helical segments or loops, each of which is given a unique identifier. Second, the 3D structure obtained at the end of the first stage of the protocol is read in to memory and all CG beads are assigned an initial coordinate in the 4^th^ spatial dimension (which we term “w”) of 0.0. Third, the geometric centers of all identified structural elements are found and the distances between all such centers are computed in 4D as the square root of x^2^ + y^2^+ z^2^+ w^2^, where x^2^, y^2^ and z^2^ are the squared separation distances along the conventional x, y, and z coordinates, respectively, and w^2^ is the squared separation distance along the added 4^th^ dimension. Structural elements are considered to be clashing or entangled if the distances between their centers are less than a specified distance (in all of the cases discussed here, this cutoff was set to 25 Å). Next, a simple Monte Carlo-like procedure is used to resolve these “clashes” by applying random displacements of up to ± 10 Å in w to all structural elements involved in clashes. It is important to note that all CG beads that are part of a single element (e.g. a double-helical stem) are assigned the same random displacement in w; in effect, therefore, each clashing element is subjected to a rigid body displacement in w. Once all elements have been assigned a random displacement, they are all checked again for clashes, and additional rounds of random displacements are made to any elements that still contain clashes until all clashes have been removed. At this point the maximum displacement in w assigned to any of the structural elements, w_max_, and the corresponding minimum displacement in w assigned to any of the structural elements, w_min_, are both recorded for later use (see below). For the largest RNAs considered here, the process of removing clashes by adding random displacements takes on the order of a few seconds; values of w_min_ and w_max_ are typically in the low hundreds of Ångstroms for the largest RNAs considered here.

At this point in the protocol, helical entanglements have been “removed” by a process that amounts to “sleight of hand”: in effect, the entanglements have been removed only by embedding the RNA into a 4D space and changing the definition of distance from one computed in 3D to one computed in 4D. The removal of entanglements in this way, however, makes it possible to begin adding in the steric interactions that were omitted from the first stage of the protocol. Steric interactions of CG beads are modelled using a purely repulsive potential function of the form E_steric_ = ε σ_bead_^12^ / r^12^ where E_steric_ is the energy of the interaction, ε is an energy parameter, σ_bead_ is the effective bead diameter, and r is the current distance between the two beads. In all the simulations described here we set ε to 1 kcal/mol, but during this second stage of the protocol σ_bead_ is increased over the course of eight short BD simulations from an initial value of 6 Å to a final value of 20 Å in intervals of 2 Å. All of these simulations are performed in 4D space with a suitably modified version of the *uiowa_bd* code, and CG beads are prevented from diffusing too far in the added spatial dimension by the addition of two half-harmonic potential functions that act along the w-coordinate. The force constant assigned to each of these “wall” functions is 1 kcal/mol/Å^2^ and their reference positions are determined by adding 10 Å to w_max_ and by subtracting 10 Å from w_min_ (see above). Each simulation is conducted for 100 ps with a timestep of 25 fs.

At the end of this second stage of the protocol, the RNA model is embedded in a 4D space and with inflated steric interactions that seek to ensure that all helical entanglements (or at least their majority; see below) have been eliminated. The goal of the third stage of the protocol is to transfer the RNA back to a 3D space. This is accomplished in a single 4D-modified *uiowa_bd* simulation in which the reference positions of the two wall functions defined above are gradually moved toward each other until they coincide. The compression of the walls progressively restricts the ability of the RNA to sample the 4^th^ spatial dimension and forces those CG bead pairs that were formerly clashing in 3D to again begin to approach each other. As they approach, however, they can seek to avoid clashes by accommodating each other’s position in a 4D space in which the 4^th^ dimension is fast disappearing; eventually, the 4^th^ dimension is removed entirely so that the CG beads are returned to 3D space. The BD simulation is conducted for a total 12.5 ns with a timestep of 6.25 fs and with the 4^th^ dimension being removed after 10 ns; a much shorter timestep is used here owing to the occasional tendency for simulations of very large RNAs to crash due to the excessive build-up of steric strain. Throughout the simulation σ_bead_ remains set to 20 Å to ensure that new helical entanglements are not allowed to form. A full description of the principles underlying this 4D simulation approach, which has applications beyond the modelling of RNA, is provided in a recent publication (36).

To verify that the RNA model is free of helical entanglements at this stage, additional code was written in-house to implement the Möller-Trumbore intersection algorithm (60) in a way similar to that already reported by the Szachniuk group (35). Briefly, the algorithm visits all loops in turn, creates triangles between the centroid of the loop and all pairs of consecutive nucleotides that are arrayed along the perimeter of the loop, and then determines if any other fragment of the RNA intersects with any of these triangles. Any intersections that are identified are first examined visually since, especially with large loop regions, many of the supposed intersections are physically plausible since they do not involve egregious entanglements of double helical regions. With most RNAs, we found that all helical entanglements were successfully removed by the second and third stages of the *uiowa_rna* protocol. In those cases where loop intersections were identified by the Möller-Trumbore algorithm, and where visual inspection indicated that they were indeed clear helical entanglements, artificial base-pairs were added to temporarily close the offending loop region and the entire protocol was repeated with the added base-pairs removed in the later stages of the protocol. While this correction process was performed manually for the RNAs presented in this study, the release version of the code includes a utility that automatically sutures all loops with temporary base-pairs that are removed in the final stages of the protocol (see Code Availability).

Once the third stage of the protocol has been completed successfully, the RNA model should be free both of steric overlaps and helical entanglements. But stage three’s use of an artificially large σ_bead_ value, which helps structural elements become disentangled from each other, also causes double-helical regions to become elongated and flattened (see third image in Figure 1). A fourth stage of the protocol, therefore, seeks to resolve this issue by performing a steepest-descent energy minimization with σ_bead_ set to a more realistic value of 10 Å, and with a number of the force constants that are applied to bonded interactions increased in magnitude so as to enforce closer adherence to an idealized double-helical A-form RNA. In particular, the force constants for all bonds are now set to 25 kcal/mol/Å; the force constants for angles are increased by a factor of 5 and the energy barriers for rotation of dihedral angles are increased by a factor of 10. A maximum step size of 0.0005 Å is allowed at each step of the energy minimization, with a maximum number of 1 million steps being carried out unless the energy on two adjacent steps changes by less than 1 × 10^-7^ kcal/mol, at which point it is terminated.

### Brownian Dynamics Simulations Including Hydrodynamic Interactions (BD-HI)

The structure obtained at the conclusion of the fourth stage of the protocol is free of steric clashes and entanglements, and has double-helical regions that, at a local level, have realistic 3D geometries (see the fourth image in Figure 1). At a global level, however, the structures of very large RNAs at this stage often have the appearance of being somewhat flat, and the relative distances between nucleotides echo their distances in the idealized 2D model that was used to start the entire process. To rid the structures of these features, therefore, a fourth and final stage of the *uiowa_rna* protocol is implemented in which a 10 µs BD-HI simulation is performed during which the RNA is allowed to freely diffuse. This simulation allows the internal dynamics of the RNA to be modelled as well as enabling the calculation of its effective hydrodynamic radius, Rh, which can then be compared with experiment (see Results). In all of the BD-HI simulations conducted here we used an energy model essentially identical to that used in our recent work (56) with the exception that bonds were added between those CG beads known to form base-pairs with each other in either the predicted secondary structure or the known 3D structure (see below). Force constants for bonds were set to 20 kcal/mol/Å, force constants for angles were set to 10 kcal/mol/rad, and, as in our recent work on proteins and RNAs (56), dihedral angles were treated as a combination of cosine functions, one with a periodicity of 1, for which the potential energy maximum was set to 0.5 kcal/mol, and one with a periodicity of 3 with the potential energy maximum set to 0.25 kcal/mol.

For BD-HI simulations of RNAs for which the 3D structure was already known (see Results), additional potential functions were added to reward the formation of tertiary contacts. This was achieved by adding Lennard-Jones potential functions between all non-bonded nucleotides for which any pair of atoms was within 5.5 Å. The well-depths of all such Lennard-Jones functions (often referred to as Gō potentials in the protein folding literature) were set to 1 kcal/mol, which is sufficiently favorable to prevent even minor unfolding events occurring during the 10 µs BD-HI simulations. For simulations aimed at describing the behavior of the same RNAs in their completely unfolded states (see Results), the favorable Lennard-Jones potential functions were omitted and all bonds between base-paired nucleotides were removed.

In all BD-HI simulations each CG bead was assigned a charge of −1 e, and their electrostatic interactions were modelled using the Debye-Hückel model with a term of the form E_int_ = 332.08 q_i_ q_j_ exp (–κ r_ij_) / (ε r_ij_), where E_int_ is the electrostatic interaction energy of two charges q_i_ and q_j_, κ is the Debye-Hückel screening parameter (related to the square root of the ionic strength, set here to 150 mM), ε is the relative dielectric constant of the solvent (set here to 78.4), and r_ij_ is the distance between the two charges. All nonbonded interactions were truncated beyond a distance of 35 Å and the list of all bead pairs interacting within this cutoff distance was recomputed at intervals of 100 simulation steps.

All of the long timescale BD-HI simulations used to model the diffusion and conformational dynamics of the RNAs include hydrodynamic interactions (HI) between CG beads, modelled at the pairwise Rotne-Prager-Yamakawa (RPY) level of theory (61,62). The radius assigned to the CG beads in the RPY calculations was optimized by comparing the hydrodynamic radii of RNAs with known 3D structures computed from BD-HI simulations with their corresponding experimental values (see Results). The optimal bead radius established from that comparison (5.5 Å) was then used in all subsequent BD-HI simulations. All simulations used a timestep of 100 fs with the RPY diffusion tensor and its Cholesky decomposition (needed for the calculation of correlated random displacements) being recomputed every 1000 simulation steps. This relatively low update frequency was used to limit the otherwise significant computational expense of the Cholesky decomposition for the largest RNAs modelled here (37), but in our simulations of five structured RNAs we found that the computed hydrodynamic radii were indistinguishable from those computed with a higher update frequency of 100 simulation steps.

All BD-HI simulations were conducted for at least 10 µs, with the first 1 µs generally discarded as an equilibration period; a typical structure obtained at the end of such a simulation is shown in the bottom image of Figure 1. For two RNA models, BD-HI simulations were extended to 100 µs in order to more fully sample the internal conformational dynamics of the molecules (see Results). For subsequent analysis, snapshots of the structure were saved at intervals of 1 ns. The computational expense of the BD-HI simulations is non-negligible. One of the largest RNAs considered in this study (the RpoB mRNA) contains 3,555 nts; a BD-HI simulation of this RNA returned approximately 1.8 µs of simulation time per day using 28 OpenMP threads on a modern compute-server containing Intel Xeon Gold 6230 CPUs.

### Analysis of BD-HI Simulations

To allow comparison with experimental hydrodynamic radii, the BD-HI simulation data were analyzed in the following way. First, the translational diffusion coefficient of the RNA was computed from analysis of the mean-squared displacement of its center of geometry during the simulation. This was achieved using the standard relationship < r^2^ > = 6 D_trans_ δt, where < r^2^ > is the mean-squared distance travelled during the observation interval δt (set here to 10 ns), and D_trans_ is the translational diffusion coefficient. The latter was then converted into an effective hydrodynamic radius, R_h_, using the Stokes-Einstein relationship, D_trans_ = k_B_T / (6πR_h_) where k_B_ is Boltzmann’s constant and T is the temperature in K (set here to 298 K). To obtain error estimates, all D_trans_ and R_h_ values were calculated separately for 3 consecutive 3 µs intervals (with the first 1 µs being omitted from consideration).

To quantify the conformational sampling during longer BD-HI simulations, the GROMACS utility `rms` was used (63). This code conducts RMSD comparisons of simulation snapshots with all snapshots best-fitted to each other. The output from this command (a .xpm file) was then read into an in-house script that converts the RMSD values from a grayscale representation to a color scale that ranges from blue to white to red. The final .xpm file was then converted into a .png format file using the image processing software GIMP (The GIMP Development Team, 2019).

To allow comparison with the cryoEM imaging results obtained by the Gelbart group (28), we calculated the shape anisotropy of structural snapshots for two of their RNAs (see below) as follows. Snapshots sampled at intervals of 1 ns, were each randomly rotated and projected onto a 2D plane 1000 times. The maximum and minimum diameter of each such projection was then calculated by rotating it around an axis running through the center of geometry of the projection and oriented perpendicular to the plane in intervals of 1°, determining the range of the projected z coordinates (i.e. the effective diameter), and storing the maximum and minimum values of this range. The anisotropy of each randomized orientation projected onto 2D was then computed as A = (M – m) / (M + M) where M and m are the maximum and minimum diameters recorded, respectively. The anisotropies computed from such an analysis for perfectly spherical, oblate, and prolate objects are expected to be 0, 0.28, and 1, respectively (28).

### Systems Studied

Four sets of RNAs have been modelled in this work. The first includes five RNAs for which experimental Rh values have been summarized by Werner (64) and whose atomic structures have either been solved experimentally, or which can be homology modelled with confidence (see below). For these RNAs, no secondary structure prediction or use of the *uiowa_rna* protocol was necessary. Instead, the atomic structures were simply converted into CG models and bonded and nonbonded parameters were assigned as described above. The structures of four of the five RNAs were downloaded directly from the RCSB (65). The fifth, that of the hepatitis delta virus (HDV) self-cleaving ribozyme, was homology modelled using as a template structure that of the near-identical construct deposited in the RCSB (RCSB ID: 4PRF, (66)). Nucleotides not found in the HDV ribozyme were removed from the RNA chain of the structure (i.e. residues 99-100, 146-159), and an atomic model of the missing P1.1 stem loop was built using the RNAComposer webserver (33) with the default settings and following input:

~~~
>HDV_stem_loop
ACAUUCCGAGGGGACCGUCCCCUCGGUAAUGG
((((((((((((((…))))))))).)))))
~~~

The resulting structure of the stem-loop was superimposed onto the trimmed 4PRF RNA chain structure using in-house code based on the superposition code PDBSUP (67). Specifically, the atoms involved in the glycosidic bonds (i.e. C1’ and either N9 for guanosine and adenosine or N1 for cytidine and uridine) of residues A1, C2, A3, U30, G31 and G32 in the stem-loop were superimposed onto the same atoms in residues A143, C144, A145, U160, G161, G162 of the trimmed 4PRF RNA structure, after which the residues A1, C2, A3, U30, G31 and G32 were removed. Finally, mutated nucleotides and the missing 3’ nucleotide of the full-length ribozyme (i.e. C85) were remodeled to produce the correct ribozyme sequence using an in-house PyMol script and the ModeRNA comparative modeling library (2).

The second set of RNAs modeled here is a set of artificial 85-nt RNAs studied by the Bartel group (68). Those authors considered three subsets of RNAs with lengths equal to that of the HDV ribozyme considered above. These subsets included: (a) RNAs with permuted sequences but with the same composition as the HDV ribozyme, (b) RNAs with essentially uniform numbers of A, C, G and U bases (with G present in one additional copy), and (c) a single poly(U)_85_ sequence containing only uridine residues that is presumed to be always unfolded. Secondary structures for each of these RNAs were predicted by RNAFold and then submitted to the *uiowa_rna* protocol. Hydrodynamic radii, R_h_, were computed from the last 9 µs of a 10 µs BD-HI simulation as described above.

The third set of 16 RNAs studied was selected from a set of large RNAs studied using fluorescence correlation spectroscopy (FCS) by a number of research teams headed by the Tuma group (69). All of the RNAs in this set were modelled using the *uiowa_rna* protocol twice: once with secondary structures predicted by RNAFold and once with secondary structures predicted by CoFold. All BD-HI simulations were conducted for 10 µs, but to assess the simulations’ ability to sample the conformational dynamics of large RNAs we continued simulations of the 961-nt BunVS RNA for a total of 100 µs. For the special cases of the ribosomal RNAs we also performed BD-HI simulations of additional models in which both the secondary and tertiary structures were taken directly from a ribosome crystal structure. To this end we conducted simulations of models of the secondary structure of both the 16S and 23S rRNA determined by the RNApdbee 2.0 webserver (70,71) from the crystal structure of the 70S ribosome (RCSB: ID 5UYM; (72)).

The final set of RNAs studied was taken from the work of the Gelbart group who used cryo-electron microscopy, small-angle X-ray scattering, and NAST-based molecular simulations to characterize ensembles of structures of large RNAs in solution (28). We selected the two RNAs (containing 975 and 1523 nts, respectively) for which anisotropy values were reported and used the protocol outlined above, together with secondary structures predicted by both RNAFold and CoFold, to create two 3D RNA models of both RNAs.

## Results

The *uiowa_rna* protocol aims to generate realistic 3D coarse-grained (CG) structures of very large RNAs that are free of steric clashes and helical entanglements. The protocol is illustrated schematically in Figure 1 for the case of the 961-nt BunVS RNA. Briefly, an initial 3D model is constructed from a 2D representation of the RNA’s predicted secondary structure, and a series of short BD simulations is then performed during which the internal parameters of the CG model (bond lengths, angles etc) are made to gradually adopt the values expected of them in 3D structures of RNAs. A novel 4D BD simulation algorithm (36) is then used to ensure that all double helical segments are disentangled from each other, steric interactions are progressively added, and a steepest-descent energy minimization is performed to allow the RNA to adopt a more geometrically realistic conformation. Finally, a 10 µs BD-HI simulation is conducted to allow the RNA to freely diffuse, to sample alternative conformations, and to allow calculation of its hydrodynamic radius, Rh, which can then be compared with experiment (see below). A movie illustrating the entire protocol on the 961-nt BunVS RNA is provided in the Supporting Information, and snapshots illustrating how, for example, loops at the ends of two nearby stem-loops become disentangled from each other during the 4D stage of the protocol are shown in Figure S1.

### Simulations of RNAs with known 3D structures

One of the end-goals of this work (see below) was to build 3D models of RNAs ranging in size up to ∼3500 nts, and use them to compute Rh values that could then be compared with corresponding experimental values (69). But since such comparisons must be based on modeled structures (for which the secondary structures are also predicted; see Materials & Methods), any differences between simulation and experiment obtained from such a comparison could, in principle, be due not only to an incorrect 3D model of the structure, but also to an incorrect description of RNA’s diffusive dynamics. One way to first separate these two potential contributions is to attempt to model the diffusive dynamics of RNAs for which *both* the 3D structure and the Rh value are already experimentally known. For a suitable test set we looked to the compilation made by Werner (64) which lists the Rh values of RNAs ranging in length from 54 to 193 nucleotides. We selected from this list five RNAs with complete or near-complete atomic structures deposited in the RCSB (73) (see Materials & Methods); these RNAs are illustrated in Figure 2A.

**Figure 2.**
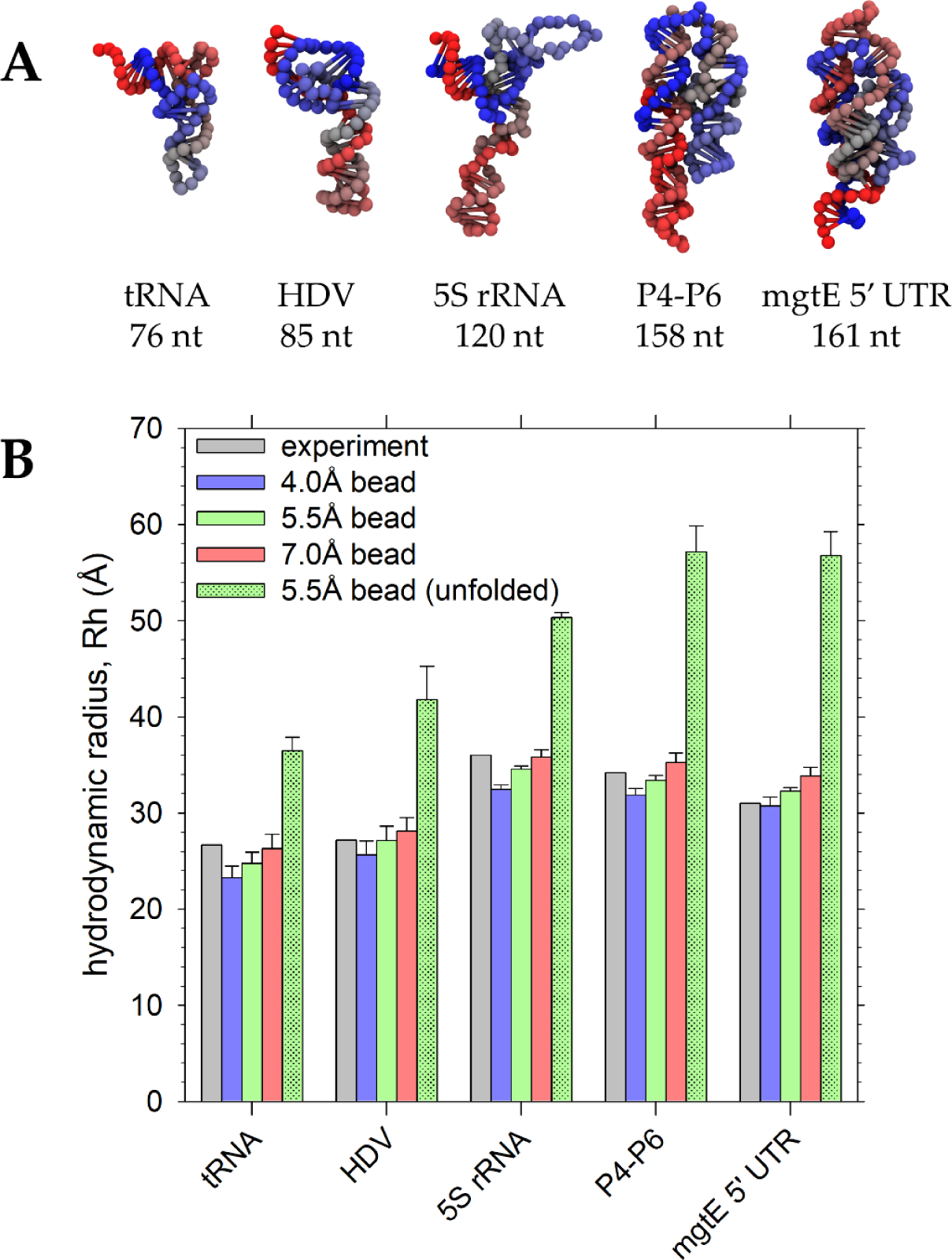
Parameterization of bead radii using RNAs with known 3D structures. **A.** Images of the RNAs used to parameterize bead radii in subsequent BD-HI simulations. **B.** Comparison of hydrodynamic radii, Rh, computed from BD-HI simulations using different candidate bead radii with experimentally reported values compiled by Werner (64). The hatched green bar shows hydrodynamic radii obtained from BD-HI simulations in which the RNA was allowed to unfold. Error bars represent the standard deviation of Rh values calculated from three contiguous blocks of the 10 µs simulation (i.e. 1.00-3.33 µs, 3.34-6.66 µs, 6.67-10.0 µs).

Using these known 3D structures as starting points, we performed 10 µs BD-HI simulations of each of the RNAs, measured their translational diffusion coefficients, and used these to calculate their effective hydrodynamic radii, Rh (see Materials & Methods). For each RNA, three independent simulations were performed, each assigning a different value to the CG beads’ hydrodynamic radii: values of 4.0, 5.5, and 7.0 Å were tested. Figure 2B compares the Rh values calculated from the BD-HI simulations with the corresponding Rh values obtained from experiment for all three values assigned to the bead radii. While all three values worked well, a bead radius of 5.5 Å produced the best overall agreement with experiment, producing an average absolute percent error of 3.6%. This bead radius was therefore used in all subsequent BD-HI simulations described in this report.

A linear regression of the simulated and experimental Rh values (Figure S2A) produces an r^2^ value of 0.91. Importantly, this is much higher than the r^2^ value obtained when the experimental Rh values are instead regressed with a power law dependence against the number of nucleotides in each RNA; the latter fit yields Rh = 8.51 N_res_^0.27^ with an r^2^ value of 0.57 (Figure S2B). This indicates that the BD-HI simulations are not simply echoing the length or the molecular weight of the RNA but are accurately capturing the influence of each RNA’s 3D shape on its diffusive dynamics. To further support the idea that the simulation-measured Rh values properly reflect the 3D structure of the RNAs, we performed additional simulations in which each of the five RNAs was allowed to unfold by eliminating all base-pairing and tertiary contacts (see Materials & Methods). In all cases, the resulting simulated Rh values (green, hatched bars in Figure 2B) become much higher than the corresponding experimental values (Figure S2C) and become in much poorer qualitative agreement with them (r^2^ = 0.57); interestingly, the ∼60% average increase in the Rh value that accompanies unfolding of the RNAs is very similar that seen in both experiments and BD-HI simulations for proteins (37). The observation that the “wrong” structure leads to incorrect Rh values suggests, therefore, that comparisons of Rh values obtained from BD-HI simulations with experimental values might be a useful way to crudely validate predicted structures of RNAs (see Discussion). Finally, it is worth noting that the simulated Rh values of the unfolded RNAs scale straightforwardly with the number of nucleotides according to a power law, with Rh = 3.57 N_res_^0.55^ (Figure S2D, r^2^= 0.98). This exponent is close to the Flory value (0.588) expected for the scaling behavior of the radius of gyration of a self-avoiding chain in a good solvent (reviewed in (74)); it is also similar to values obtained from regressing experimental Rh values of structured RNAs by Werner (64).

### Simulations of 85 nt RNAs with unknown 3D structures

A set of experimental data that provide the first test of the *uiowa_rna* protocol’s ability to generate reasonable 3D models of RNAs comes from the Bartel group who, in addition to reporting the Rh value for the hepatitis delta virus (HDV) ribozyme appearing in Figure 2, also reported Rh values for 21 synthetic RNAs, each containing 85 nts (68). While each of the 21 synthetic RNAs has a unique sequence, they can be separated into one of three different classes. The first class, containing ten RNAs, is a so-called permuted (P) class, in which all sequences have the same nucleotide content as found in the HDV ribozyme; the nucleotide content of these sequences is ∼71 % G or C nts. The second class, also containing ten RNAs, is the so-called isoheteropolymer (I) class, in which each type of nucleotide is present at an effective ratio of 1:1:1:1 (with G overrepresented by a single nucleotide in all cases). The remaining class, containing only a single RNA, is a 85-nt poly-uridine, i.e. poly(U)_85_; in contrast to the other RNAs, this RNA is unable to form base pairs in solution and, as a result, has a drastically increased experimental Rh value compared to its P class and I class counterparts (68). This dataset provides the first stringent test of our protocol because the experimental secondary and tertiary structures of the 21 RNAs are not known with certainty, although their secondary structures can probably be predicted with reasonable accuracy.

We used our protocol, therefore, to create 3D models for all 21 RNAs in the Bartel group’s data set and subjected each of them to a 10 µs BD-HI simulation using the optimized bead radius of 5.5 Å. The predicted secondary structures (all predicted using RNAFold; see Materials & Methods) and the tertiary structures at the end of the 10 µs BD-HI simulation period are shown in Figure 3A. The average Rh values calculated for each class are illustrated in Figure 3B; since our protocol builds 3D models solely on the basis of a predicted secondary structure (i.e. it does not, at this stage, include additional tertiary contacts; see Discussion), we compare our results with similarly averaged Rh values reported by the Bartel group obtained in the absence of Mg^2+^ but in the presence of 30 mM KCl. While the experimental Rh values for the P and I class RNAs are very similar to each other, the mean Rh value of the more GC-rich P class sequences is lower at 31.3 Å, compared to 32.7 Å for the I class (see the gray bars in Figure 3B). According to a two-tailed T-test, the difference between these two mean experimental values, while small, is significant at the level p=2.28 × 10^−4^. Encouragingly, the class-averaged Rh values obtained from the simulations match the experimental Rh values rather well, with mean values of 31.2 Å and 32.1 Å for the P and I class respectively (see the yellow bars in Figure 3B); in this case, however, the difference between the two mean values is significant only at the level p = 0.057. Within each class, the trends for the Rh values are not reproduced by our models, but it is probable that, for both the experimental and simulated datasets, the Rh values of all RNAs in each class are within error of each other: no error estimates were provided for the experimental values (68), but the error estimates for individual simulations reported here are typically ∼1.5 Å. Finally, the much higher Rh value for poly(U)_85_ is, as expected, qualitatively reproduced by the BD-HI simulations, although it is somewhat underestimated at 40.5 Å relative to the experimental value of 45.5 Å. Overall, then, the simulation models presented here reproduce the Bartel group’s data quite well (see Figure 3B), with the Rh values of the HDV ribozyme, the P and I class variants, and the poly(U)_85_ sequence all qualitatively reproduced, and, with the exception of the latter, quantitatively reproduced. In addition, these results again demonstrate that the simulations are appropriately sensitive to the structure details of the 3D models. Individual Rh values for all RNAs in this data set are reported in Table S1.

**Figure 3.**
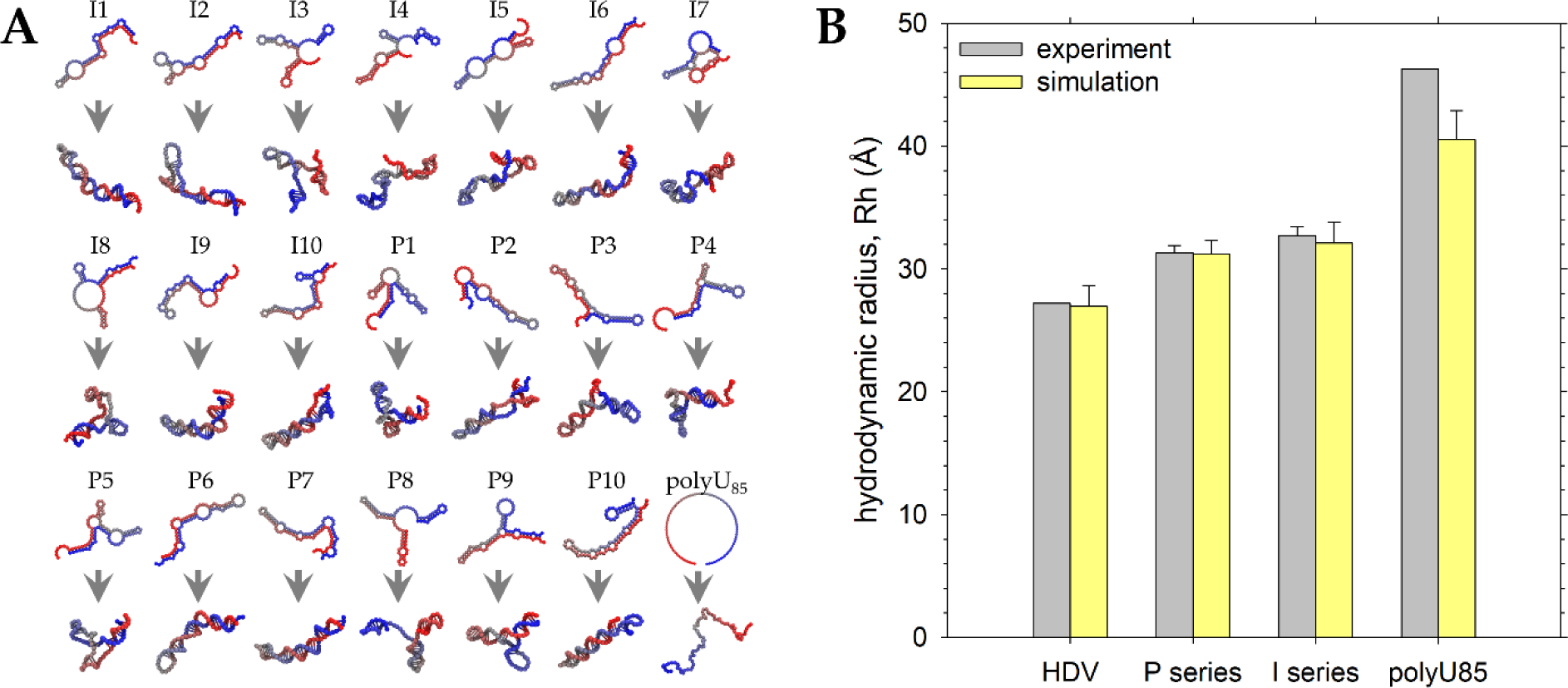
Application of the 3D modeling protocol to 21 RNAs of length 85 nts. **A.** Images showing the predicted secondary structures of the 21 RNAs in the Bartel group’s dataset (top row in each pair of images), and the resulting 3D models at the end of a 10 µs BD-HI simulation (bottom row). **B.** Comparison of mean hydrodynamic radii, Rh, of the 10 P-class RNAs, the 10 I-class RNAs, and the U_85_ RNA with their corresponding experimentally reported values (68); also shown are simulated and experimental results for the native state of the 85-nt HDV ribozyme. Error bars on the P-class and I-class RNA data represent the standard deviation of the mean Rh values calculated for the 10 RNAs in each class; the error bars on the simulated data for the HDV and U_85_ RNAs were calculated as in Figure 2; error bars on the experimental data for the HDV and U_85_ RNAs were not available.

### Simulations of large RNAs – comparison with experimental Rh values

Having shown that good results can be obtained for smaller RNAs whose 3D structures are not explicitly known but whose secondary structures can be predicted with some degree of confidence, we built 3D models of 16 much larger RNAs and attempted to validate them by comparing their simulated Rh values with corresponding experimental values obtained from fluorescence correlation spectroscopy (FCS) measurements by a number of research teams led by the Tuma group (69). Since our protocol hinges on having available a reliable secondary structure prediction, and since these RNAs are sufficiently large that there is likely to be considerable uncertainty in their secondary structures, we repeated the entire analysis using secondary structures predicted by both RNAFold and CoFold.

Figure 4A shows snapshots of four representative RNAs with their 3D models built using minimum free energy (MFE) secondary structures predicted by RNAFold (48); snapshots of all 16 models are shown in Figure S3. The structures vary significantly in terms of their compactness, but one notable aspect is that, in most cases, the 5’ and 3’ ends are near each other owing to base-pairing interactions between them; this is a known feature of methods such as RNAFold (75). Figure 4B shows snapshots of the same four RNAs with their 3D models built using MFE secondary structures predicted by CoFold (49), a method that attempts to implicitly account for the impact that cotranscriptional folding can have on secondary structure formation. The structures predicted using CoFold are generally less compact, more string-like, and (as expected) show much less evidence of long-range base-pairing between the 5’ and 3’ ends; snapshots of all models built using CoFold-predicted secondary structures are shown in Figure S4.

**Figure 4.**
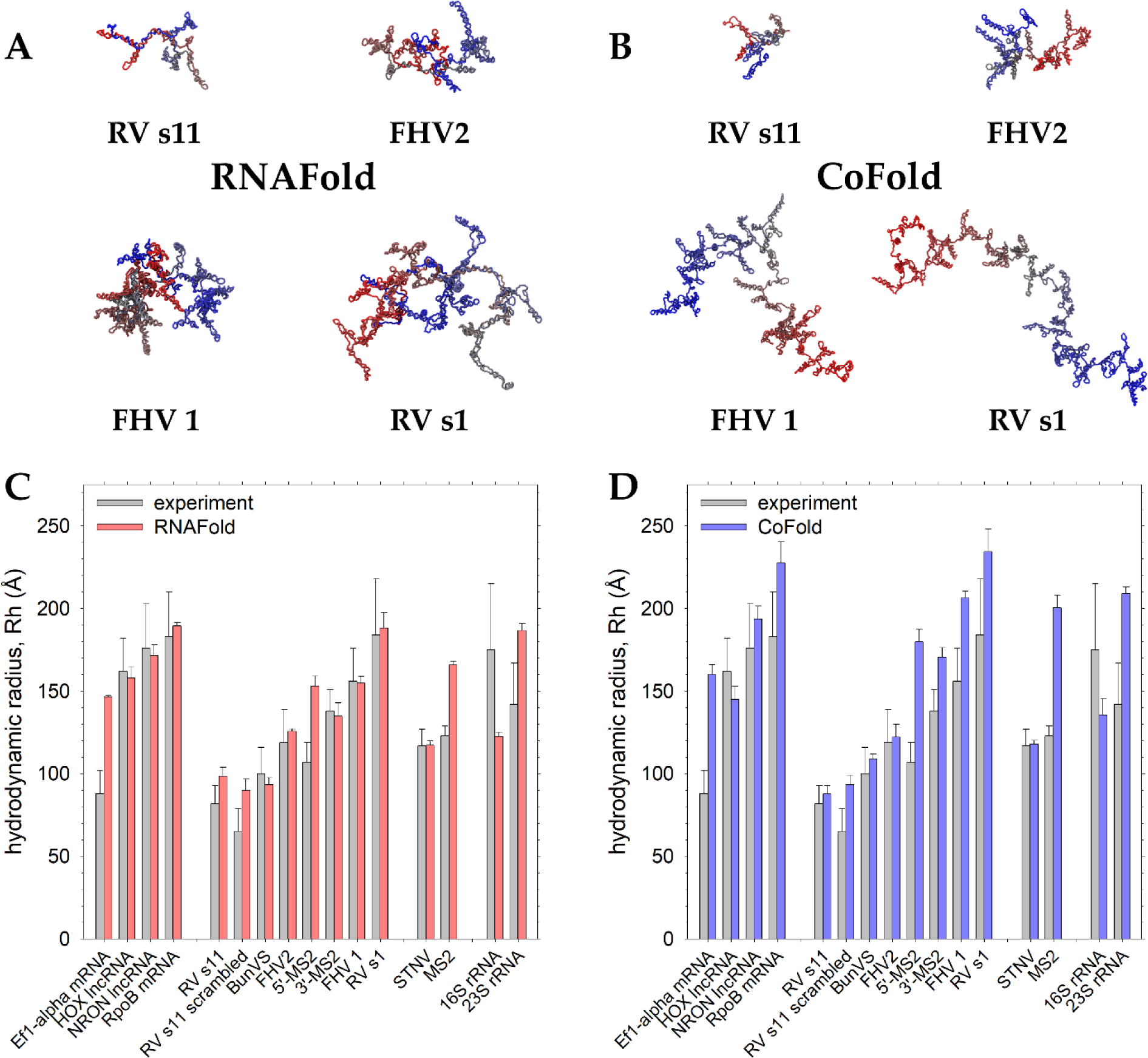
Application of the 3D modeling protocol to 16 RNAs of length up to ∼3500 nts. **A.** Representative images of 3D models built using RNAFold-predicted secondary structures at the end of a 10 µs BD-HI simulation. **B.** Same as A but showing models built using CoFold-predicted secondary structures. **C.** Comparison of mean hydrodynamic radii, Rh, of 16 RNAs selected from the dataset reported by Tuma and co-workers (69), calculated from BD-HI simulations of 3D models built using RNAFold-predicted secondary structures. **D.** Same as C but for 3D models built using CoFold-predicted secondary structures. Error bars on the simulated data were calculated as in Figure 2; error bars on the experimental data were taken from Table 1 of Borodavka et al. (69).

Figure 4C compares the Rh values obtained from BD-HI simulations with experiment for models built using secondary structures predicted by RNAFold. Here we separate the RNAs into four classes: (a) lncRNAs and mRNAs, (b) fragments of viral genomes, (c) complete viral genomes, and (d) ribosomal RNAs. The level of agreement between simulation and experiment differs sharply between classes. In the case of the lncRNAs and mRNAs, the agreement between the RNAFold-predicted structures and experiment is rather good, with the sole exception of the Ef1-α mRNA, for which the experimental value is known to be unusually low (69). In the case of the fragments of viral genomes, the agreement between the RNAFold-predicted structures and experiment is again good, with the most conspicuous outlier being the 5’-MS2 fragment, for which the experimental value has again been noted to be unusually low relative to other genomic fragments (69). The RNAFold-predicted Rh values for the unscrambled and scrambled RV s11 fragments are also on the high side, but they at least qualitatively reproduce the experimental result that the variant with the scrambled sequence has the smaller Rh value. For the complete viral genomes, the prediction for STNV is excellent, while the prediction for MS2 is much too high; the latter result echoes the earlier result for the 5’-MS2 fragment and is consistent with the predicted secondary structure of the latter being largely retained in the full-length genome prediction. Finally, for the ribosomal subunits, the agreement between simulation and experiment is abysmal: the Rh value for the 16S rRNA is hugely underestimated, while that of the 23S rRNA is hugely overestimated. Possible reasons for these clear failures are considered in a subsequent section.

Figure 4D compares the Rh values obtained from BD-HI simulations with experiment for models built using secondary structures predicted by CoFold. As expected, given the snapshots shown in Figure 4B, the Rh values obtained from CoFold-predicted structures are generally higher than those obtained with RNAFold (see Figure S5A for a comparison). They also appear to be in slightly poorer agreement with experiment, but the same sets of RNAs are identified as outliers: relative to experiment, the predicted Rh values are overestimated for Ef1-α mRNA, 5’-MS2, full-length MS2, and the 23S rRNA, and drastically underestimated for the 16S rRNA. We note that for both RNAFold and CoFold-derived simulations, BD-HI simulations of 10 µs duration appears to be sufficient to obtain converged estimates of the Rh values since the Rh values obtained from 3 contiguous 3 µs blocks are, for all of the 16 large RNAs studied here, very similar to each other (Figure S6).

Before considering possible causes of the very poor results obtained with the rRNAs, it is first worth examining a few other trends in the predicted Rh values. First, as was the case with the unstructured RNAs (see above), both sets of predicted Rh values fit well to a power law of the type proposed by Werner (64): for RNAFold, we obtain Rh = 6.14 N_res_^0.414^ (Figure S5B, r^2^= 0.87), for CoFold, we obtain Rh = 2.07 N_res_^0.571^ (Figure S5C, r^2^= 0. 94). The higher exponent achieved with the CoFold-predicted structures is consistent with their generally more string-like structures. Second, the observation that the CoFold-predicted Rh values can differ substantially from those obtained using RNAFold indicates that the simulated Rh values have a measurable sensitivity to the method used to predict the secondary structure. To test the extent to which alternative, thermally accessible secondary structures predicted with the same method might also lead to different Rh values we used RNAFold to make five alternative models of two of these RNAs: the 961-nt BunVS and the 1400-nt FHV2 RNAs (see Materials & Methods). Figure S7A depicts the predicted secondary structures of the BunVS variants together with their corresponding 3D models; Figure S7B shows the same but for the FHV2 variants. As expected, for both RNAs, the MFE structures contain greater numbers of base-pairs than the non-MFE variants: 63.3% of nucleotides are base-paired in the FHV2 MFE, for example, while its non-MFE variants have a mean of 60.7% of the nucleotides base-paired. While many of the MFE’s base-pairs are retained in the variants, there are significant differences with ∼21% of nucleotides paired differently from the MFE in the BunVS variants and ∼27% of nucleotides paired differently in the FHV2 variants. Despite these differences, the resulting Rh values of all variants are very similar to those of the corresponding MFEs (Figure S7C). This indicates that while the Rh values obtained from the BD-HI simulations can be clearly sensitive to the overall 3D shape of the RNA (see, e.g. poly(U)_85_), and to large differences in predicted secondary structures (e.g. RNAFold versus CoFold), they appear to be less sensitive to the fine details of the secondary structure predicted by a given method.

### Alternative models for ribosomal RNAs

In the above comparisons, two of the most notable outliers are the 16S and 23S rRNAs: with both RNAFold and CoFold, the predicted 16S rRNA models give Rh values that drastically underestimate the reported experimental value of 175 ± 40 Å, while the predicted 23S rRNA models give Rh values that significantly overestimate the experimental value of 142 ± 25 Å. The availability of atomic structures of the ribosome allows us to consider the use of alternative 3D models that, together with the known secondary structure and/or tertiary contacts, are taken directly from the crystal structure instead of being predicted (see Materials & Methods).

For the 16S rRNA, a BD-HI simulation of a model that uses the secondary structure seen in the 5UYM ribosome crystal structure (yellow bar in Figure 5) gives an Rh value that is lower than the values obtained with the *uiowa_rna*-predicted models, thereby pushing the computed value even further away from the reported experimental value. As would be anticipated, a BD-HI simulation of a model that adds stabilizing tertiary contacts in addition to the experimental secondary structure (see Materials & Methods) gives an Rh value that is even lower (green bar in Figure 5). Given that we showed earlier that BD-HI simulations accurately reproduce Rh values for RNAs whose 3D structure is known, the latter result suggests that the experimental Rh value for the 16S rRNA is fundamentally incompatible with any of the predicted structures. In fact, the closest that we can come to reproducing the experimental values is by using the *uiowa_rna* protocol to build a 3D model of a predicted dimer of 16S rRNAs: the secondary structure for this hypothetical dimeric form, which was predicted using RNAcofold ((53); see Materials & Methods), is shown in Figure S8. Even then, however, the Rh value obtained from a BD-HI simulation of this dimeric form (pink bar in Figure 5) remains some way short of the reported experimental value.

**Figure 5.**
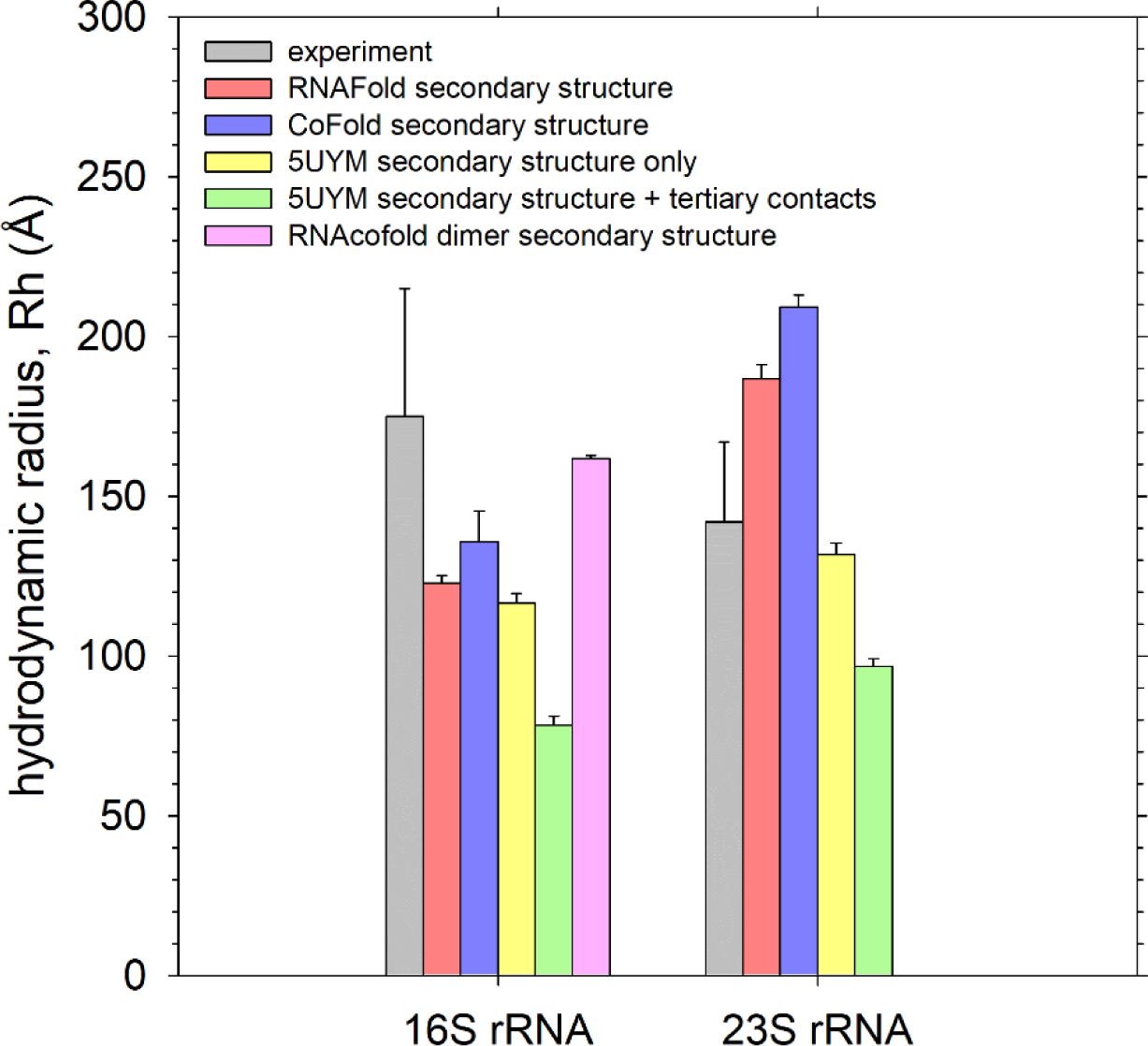
Effects of different secondary structures on simulated Rh value of ribosomal RNAs. Comparison of mean hydrodynamic radii, Rh, of 16S and 23S rRNAs calculated from BD-HI simulations of 3D models built starting from different secondary structures. Error bars on the simulated data were calculated as in Figure 2; error bars on the experimental data were taken from Table 1 of Borodavka et al. (69).

In contrast to what was seen with the 16S rRNA, use of a 3D model that starts with the secondary structure seen in the ribosome crystal structure for the 23S rRNA results in a computed Rh value that is in much better agreement with experiment (yellow bar in Figure 5). Again, adding in stabilizing tertiary contacts causes the Rh value to drop further and to deviate more, therefore, from the experimental value (green bar in Figure 5). While we have no explanation for the poor results obtained with the 16S rRNA, there does at least appear to be a straightforward explanation for the poor results obtained for the 23S rRNA: this is that both the RNAFold and CoFold predicted secondary structures are likely to be a long way from correct.

### Simulations of large RNAs – comparison with cryoEM images

While our primary focus here is on comparison with hydrodynamic data for large RNAs, it is also of interest to compare 3D models built using the present protocol with structures of two large RNAs directly imaged by the Gelbart group using cryoelectron microscopy (28). As with the 16 large RNAs considered above, we built 3D models of both RNAs using secondary structures predicted with both RNAFold and CoFold. To facilitate the comparison, we analyze the predicted structures using methods that closely track those used experimentally; in particular, we calculate the shape anisotropy of conformations sampled during the simulations to determine the extent to which their overall shapes can be considered oblate, prolate (or scalene) ellipsoids (see Materials & Methods). The maximum and minimum diameters obtained from projections of the structures are plotted versus their anisotropies in Figure S9; each is in good qualitative agreement with the corresponding experimental plots shown in Figure 2 of Gopal et al. (28). Plots of the anisotropy as a cumulative average over the course of the BD-HI simulation trajectories are shown in Figure S10; these indicate that, in all cases, the values typically drop substantially during the first 1 µs of the simulation (confirming again that this period should be considered an equilibration period), but begin to converge after that. Using snapshots sampled from the last 9 µs of the 10 µs BD-HI simulations, we obtain mean anisotropy values of 0.21 and 0.23 for the 975- xand 1523-nt RNAs using RNAFold-predicted secondary structures, and 0.25 and 0.24 using CoFold-predicted secondary structures, respectively. These values are somewhat lower than the values reported by the Gelbart group (28), who reported values of 0.32 and 0.34, respectively, which suggests, perhaps surprisingly, that that our predicted models are somewhat more globular than is suggested experimentally. Using the calculated anisotropy values and Figure 3 of the Gelbart group’s work (28), we can estimate that our models behave similar to ellipsoids with aspect ratios for the principal axes of 3.9:3.9:1 and 4.1:3.9:1 using RNAFold, and aspect ratios of 4.1:3.5:1 and 3.2:2.5:1 using CoFold. These ratios compare quite well with the somewhat more prolate set of values of reported by the Gelbart group of 4.2:2.4:1 and 5.3:3.0:1, respectively (28).

To expand on the exploration of shape anisotropies, we performed similar analyses for the 16 large RNA models discussed earlier, the results of which are listed in Tables S2 and S3. In all cases, the mean anisotropy values calculated for the models predicted by *uiowa_rna* are somewhat lower than those reported for the two RNAs studied by the Gelbart group: the mean anisotropy values for all 16 RNAs are 0.23 ± 0.05 and 0.27 ± 0.05 for the RNAFold-derived and CoFold-derived models, respectively. Importantly, however, the principal axes ratios for the corresponding ellipsoid models are again quite similar to those derived by Gelbart group – averaging 3.4:2.8:1 and 4.0:3.0:1 for RNAFold- and CoFold-derived models, respectively, thereby supporting their interesting suggestion that all large RNAs are likely to have similar overall shapes (28).

### Simulations of large RNAs – conformational flexibility

Finally, we consider an additional feature that comes “for free” from BD-HI simulations that are ostensibly intended to model overall diffusive dynamics: these are insights into the internal conformational dynamics of the RNAs. Visual inspection of the 10 µs BD-HI simulation trajectories of the 16 large RNAs indicates that all models exhibit a great deal of conformational flexibility, which is perhaps to be expected given that the models contain only secondary structure and no tertiary contacts. But to expand on this impression, we performed 100 µs BD-HI simulations of the 961nt BunVS RNA as a representative example. Figure 6A shows an overlay of 11 superimposed snapshots, sampled at intervals of 1 µs, from the BD-HI simulation performed with a RNAFold-predicted secondary structure; Figure 6B does the same but from a BD-HI simulation performed with a CoFold-predicted secondary structure. It can be seen that large-scale conformational fluctuations occur in both simulations, so it is not a surprise to find that pairwise comparisons of superimposed snapshots produce often large RMSD values. Figure 6C presents these RMSDs in 2D form for the RNAFold-based simulation, while Figure 6D does the same for the CoFold-based simulation. In both plots, blue areas indicate pairs of conformations that are very similar to each other, while red areas indicate pairs of conformations that are very different from each other. Despite the fact that very large-scale conformational fluctuations frequently occur, the fact that blue areas are found at positions a long way from the diagonal indicates that similar conformations are also repeatedly sampled at very different timepoints in both simulations. This suggests that while the internal conformational landscapes of large RNAs are undoubtedly broad (and would become even more so if allowed to sampled alternative secondary structures), BD-HI simulations performed on a timescale of 100 µs are likely to cover a significant fraction of the space.

**Figure 6.**
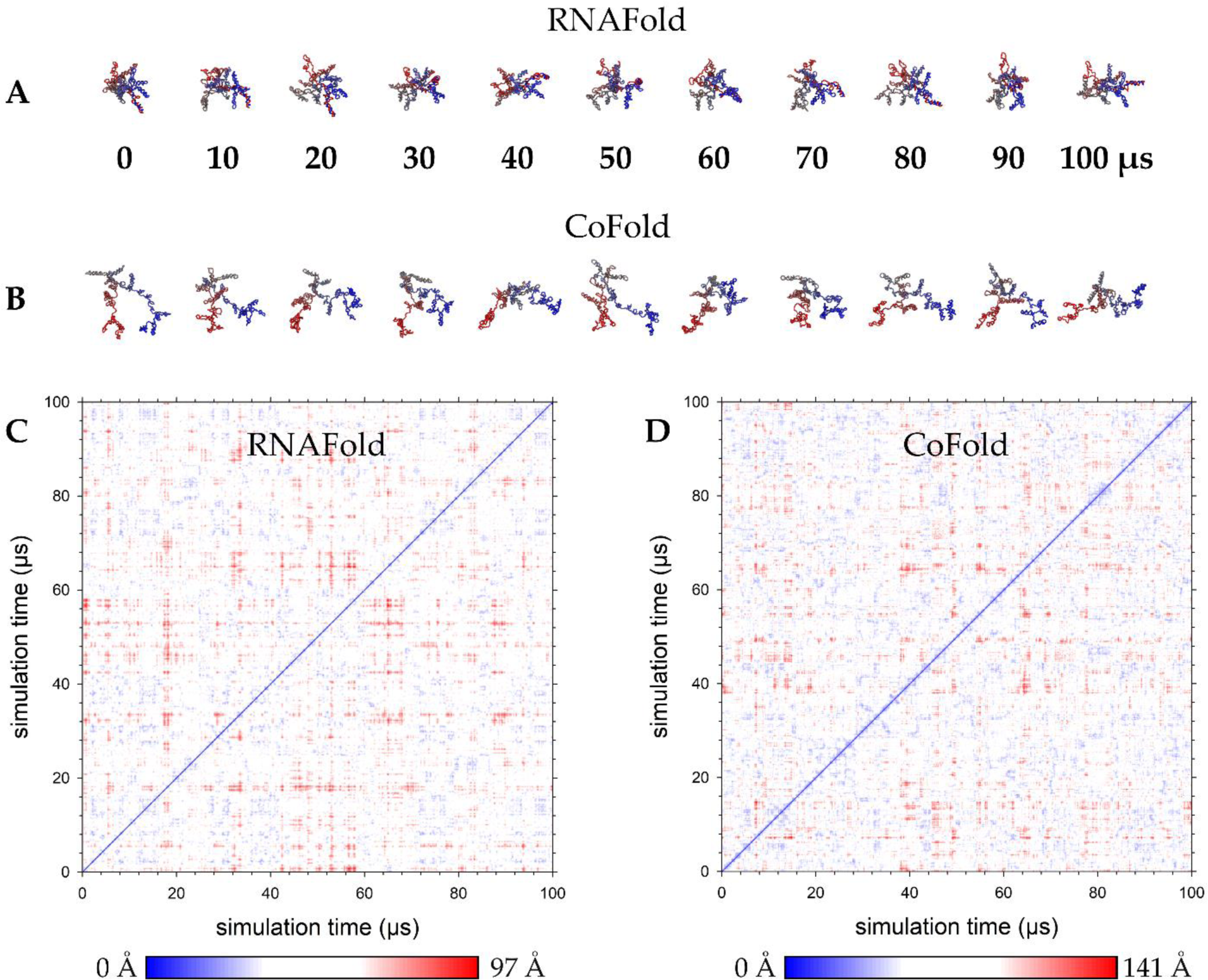
Conformational sampling during 100 µs BD-HI simulations of the 961-nt BunVS RNA. **A.** Images showing conformations sampled from a BD-HI simulation of a 3D model built using an RNAFold-predicted secondary structure; snapshots were sampled at intervals of 10 µs and RMSD-aligned to the energy-minimized 3D model using VMD. **B.** Same as **A** but from a BD-HI simulation of a 3D model built using a CoFold-predicted secondary structure. **C.** 2D map plotting the RMSD of structures sampled throughout the 100 µs BD-HI simulation of a 3D model built using an RNAFold-predicted secondary structure. Blue regions represent similar pairs of conformations; red regions indicate highly dissimilar pairs of conformations. **D.** Same as **C** but for a 3D model built using a CoFold-predicted secondary structure.

## Discussion

There are two principal results of the work reported here. First, we have described a computational protocol that is capable of building CG models of large RNAs and we have demonstrated its use here with RNAs of lengths up to ∼3500 nts. One important feature of the protocol is its use of an idealized representation of the RNA’s predicted secondary structure as its initial 3D structure. This step ensures that the model already contains all of the base-pairing interactions specified by the secondary structure and obviates, therefore, the need for effort to be expended in bringing very distant nucleotides into contact, which would be the case if a linear or circular initial model was instead used. Another significant feature of the protocol is its use of a novel 4D simulation approach (36) that allows steric interactions to be easily introduced and that, crucially, allows double-helical segments to be disentangled from each other, in a way that is essentially automatic. This ability to remove non-physical structures (35) that are often built (accidentally) into large RNA models by other methods is likely to facilitate conversion of the CG models presented here into atomic models using methods, such as C2A (76), that are already available. More benign entanglements can still occur – such as those involving large loops that are marginally encroached upon by hairpins – but these can, if desired, be resolved manually or, as is available in the release version of the *uiowa_rna* code, using an automated procedure that temporarily sutures large loops with additional bonds (base-pairs) that are later removed (see Materials & Methods).

The second major outcome of the present work is the demonstration that BD-HI simulations, while somewhat computationally expensive to perform, can be applied to even rather large RNAs, thereby simultaneously providing information on both the molecule’s diffusive and conformational dynamics. Application of the same method to even larger RNAs is becoming increasingly feasible with continued efforts to accelerate BD-HI simulations (56,77). In terms of the specific simulation methodology used here, the clearest antecedent of the present work is a report from the Garcia de la Torre group in which BD-HI simulations were used for the first time to model two RNAs (38). The inclusion of HIs is important for implicit solvent simulation methods such as BD and Langevin dynamics to correctly reproduce translational and rotational diffusion: in the absence of HIs, for example, the simulated translational diffusion coefficients of proteins can be underestimated by more than an order of magnitude (37). The excellent agreement with experimental Rh values obtained here for RNAs with known atomic structures (Figure 2), together with the Garcia de la Torre group’s demonstration (38) that BD-HI simulations of CG RNA models also match transport properties predicted by the sophisticated hydrodynamic modelling program HYDROPRO (78), have at least one important implication. They indicate that BD-HI simulations can be used to crudely validate a modelled structure if a corresponding experimental Rh value is available. There are, of course, limitations to what can be concluded about the correctness of a 3D model: we have shown here, for example, that five different predicted secondary structures all result in Rh values that are essentially identical (Figure S7). This indicates that agreement between simulated and experimental Rh values only signals that the overall dimensions of the molecule are correct: it does not provide any information on the likely correctness of the underlying secondary structure. That said, while a correctly reproduced Rh value does not guarantee that the details of the 3D model are correct, the corollary is probably true: an incorrectly reproduced Rh value almost certainly indicates that the 3D model is incorrect (see below).

While the above highlights the advantages of the BD-HI method, the application of the method to the large RNAs studied by the Tuma group and collaborators (69) hints at its inherent limitations. It is probably fair to say that the level of agreement that we obtain with experiment is reasonable but underwhelming, and it is important to identify the causes of the more drastic discrepancies. Leaving to one side, for now, the ribosomal RNAs, four of the RNAs have their Rh values consistently overestimated by the *uiowa_rna* protocol, regardless of whether RNAFold or CoFold is used to predict the secondary structure. These are: RV s11, Ef1α, full-length MS2, and its 5’-domain (5’-MS2), whose predicted secondary structure is almost 100% preserved in the predicted secondary structure of the full-length genome. A possible explanation for the errors with these RNAs, based in part on some of the issues discussed in the Tuma group’s work (69), can be proposed as follows. One notable aspect of the experimental dataset is that most of the RNAs exhibit a clear compaction when the ionic conditions of the experiment are changed from low salt conditions to 20 mM Mg^2+^: these changes are consistent with the formation of salt-dependent tertiary interactions that are stabilized by the added Mg^2+^ ions. Strikingly, however, the four RNAs for which our Rh values are consistently overestimated do not show this trend, but instead exhibit much smaller changes (<10% decreases) to their Rh values when 20 mM Mg^2+^ is added. A salt-independent Rh value could, in principle, be interpreted in one of two ways: it could either imply that there are no tertiary interactions in these RNAs under any salt conditions, or it could imply that such tertiary interactions persist even at very low salt concentrations. The low-salt experimental Rh values of all four of these RNAs are lower than would be expected on the basis of their molecular weights, and they were explicitly identified as outliers for this reason by Borodavka et al. (69). The most likely interpretation, therefore, is that these RNAs have tertiary interactions that are stable even under low salt conditions (69). Since tertiary interactions are not explicitly accounted for in the present protocol, the overestimation of the Rh values for these RNAs is not surprising. The remaining questions for these RNAs, then, are perhaps more biological than biophysical, namely, why do these RNAs behave the way they do experimentally? Some interesting potential explanations have already been suggested by Borodavka et al. (69).

From a purely biophysical perspective, the most puzzling results obtained here are those for the two ribosomal RNAs. Given that, in contrast to many of the other RNAs in the experimental dataset, both of these RNAs have evolved to specifically bind a host of proteins it is perhaps no surprise to find that accurate prediction of their structures and/or hydrodynamic properties presents the biggest obstacle to the RNA-only protocol reported here. But a simple examination of the relative experimental values for the two RNAs also indicates that their behavior is likely to be highly non-trivial to reproduce. The 16S rRNA contains 1542 nucleotides and, in the Borodavka et al. study, has a reported experimental Rh of 175 ± 40 Å in low salt conditions; the 23S rRNA, on the other hand, has 2904 nucleotides, i.e. nearly double the number of nucleotides of the 16S, and yet has a smaller reported Rh of 142 ± 25 Å. This qualitative reversal in the ordering of the Rh values persists in the experimental data obtained in 20 mM Mg^2+^ (69). While the Rh values for both RNAs have been shown to be sensitive to the addition of Mg^2+^ and thus, by the argument that we advance above, should be possible for us to predict with some degree of accuracy, there is no question that our results are poor for both RNAs.

The simpler of the two cases to explain seems to be that of the 23S rRNA. Its Rh value is over-estimated by our models, regardless of whether we use secondary structures predicted by either RNAFold or CoFold and in this sense, therefore, it represents a clear failure of our protocol. When we instead use the secondary structure evident in the crystal structure of the 70S ribosome, the Rh value is nicely reproduced (see above). Given the known stabilizing role of ribosomal proteins, use of the crystal structure’s secondary structure in a simulation of the RNA alone is unlikely to be completely appropriate. Nevertheless, it appears to be a better model of the protein-free RNA than that predicted by either RNAFold or CoFold. Notably, when a further simulation is performed in which the crystal structure’s tertiary interactions are preserved, the resulting Rh value underestimates the experimental value very significantly. This result is to be expected given that, in reality, the tertiary folding of the 23S rRNA is contingent upon the binding of ribosomal proteins that are absent from both the experiments and the simulations.

The case of the 16S rRNA is more difficult to explain. In particular, this is the only large RNA for which the simulated Rh values substantially underestimate the reported experimental value. It is difficult to avoid the conclusion that the structure of the 16S rRNA alone in low-salt conditions must somehow be vastly different both from that seen in the ribosome crystal structure and from the models predicted by RNAFold or CoFold. The large reported uncertainty in the experimental Rh value (± 40 Å) is noteworthy, but a search of the literature does find support for the mean value of 175 Å reported by Borodavka et al. (69): a value of 176 Å, for example, was reported for the radius of gyration based on SAXS measurements (79), and an approximate value of 175 Å can be read for the Rh value derived from both DLS and sedimentation measurements in Figure 1 of Timchenko et al. (80). But lower values have been described in other studies, with the reported values depending significantly on the details of the salt composition of the buffer and on the temperature. For example, a Rh value of 125 Å has been reported in the absence of Mg^2+^ at 4°C (81) based on sedimentation data reported elsewhere (82), and a Rh value of 154 Å was reported based on viscosity data in a Mg^2+^-free buffer (83). Interestingly, at least two of these studies noted a tendency for the 16S rRNA to dimerize, albeit only in higher salt conditions (81,83); against this, however, it is also to be noted that the FCS experiments described by the Tuma group were carried out at very low RNA concentrations with the specific purpose of minimizing the potential for aggregation such as this to occur (69). It is therefore intriguing that when we use our protocol to predict the structure of a 16S rRNA dimer (whose secondary structure was predicted by RNAcofold) we obtain a Rh value that is much closer to the reported experimental value (see pink bar in Figure 5).

If we consider only those non-ribosomal RNAs whose Rh values have been shown to be salt-dependent, we obtain an excellent correspondence between the predicted Rh values using RNAFold-predicted secondary structures and experiment (see Figure S11). This suggests that the modelling protocol outlined here basically works, at least for those RNAs that we hypothesize do not contain any low-salt tertiary interactions. But, for more recalcitrant cases that have significant tertiary interactions, it is likely that Rh values predicted by the current protocol will be substantially overestimated. For such RNAs, better results are likely to be obtained using a secondary structure that is either constrained by experimental data, or that is predicted with advanced deep learning methods that have improved abilities to predict long-range tertiary interactions (84,85). Extending the current protocol to include the latter types of interactions – which in principle could be dealt with by including additional distance-dependent potential functions – is an obvious direction for future work.

## AVAILABILITY

All computer code necessary to run the simulations described here will be made available to reviewers at the time of manuscript review. Upon acceptance of the manuscript for publication, the computer code will be available to the community at the following GitHub repository (https://github.com/Elcock-Lab/uiowa_rna)

## ACKNOWLEDGEMENT

This research was supported in part through computational resources provided by The University of Iowa, Iowa City, Iowa.

## REFERENCES

1. Frellsen, J., Moltke, I., Thiim, M., Mardia, K.V., Ferkinghoff-Borg, J. and Hamelryck, T. (2009) A Probabilistic Model of RNA Conformational Space. PLOS Computational Biology, 5, e1000406.

2. Rother, M., Rother, K., Puton, T. and Bujnicki, J.M. (2011) ModeRNA: a tool for comparative modeling of RNA 3D structure. Nucleic acids research, 39, 4007–4022.

3. Sharma, S., Ding, F. and Dokholyan, N. (2008) iFoldRNA: Three-dimensional RNA Structure Prediction and Folding. Bioinformatics (Oxford, England), 24, 1951–1952.

4. Boniecki, M.J., Lach, G., Dawson, W.K., Tomala, K., Lukasz, P., Soltysinski, T., Rother, K.M. and Bujnicki, J.M. (2016) SimRNA: a coarse-grained method for RNA folding simulations and 3D structure prediction. Nucleic acids research, 44, e63–e63.

5. Parisien, M. and Major, F. (2008) The MC-Fold and MC-Sym pipeline infers RNA structure from sequence data. Nature, 452, 51–55.

6. Wang, J., Wang, J., Huang, Y. and Xiao, Y. (2019) 3dRNA v2.0: An Updated Web Server for RNA 3D Structure Prediction. Int J Mol Sci, 20, 4116.

7. Das, R. and Baker, D. (2007) Automated *de novo* prediction of native-like RNA tertiary structures. Proceedings of the National Academy of Sciences, 104, 14664.

8. Watkins, A.M., Rangan, R. and Das, R. (2020) FARFAR2: Improved De Novo Rosetta Prediction of Complex Global RNA Folds. Structure, 28, 963–976.e966.

9. Zhang, D., Li, J. and Chen, S.-J. (2021) IsRNA1: De Novo Prediction and Blind Screening of RNA 3D Structures. Journal of Chemical Theory and Computation, 17, 1842–1857.

10. Chenjie, F., Wenkai, W., Renmin, H., Ziyi, W., Lisa, Y., Zongyang, D., Hong, W., Fa, Z., Zhenling, P. and Jianyi, Y. (2022) Accurate *de novo* prediction of RNA 3D structure with transformer network. bioRxiv, 2022.2010.2024.513506.

11. Robin, P., Gilbert, S.O. and Yang, Z. (2022) *De Novo* RNA Tertiary Structure Prediction at Atomic Resolution Using Geometric Potentials from Deep Learning. bioRxiv, 2022.2005.2015.491755.

12. Baek, M., McHugh, R., Anishchenko, I., Baker, D. and DiMaio, F. (2022) Accurate prediction of nucleic acid and protein-nucleic acid complexes using RoseTTAFoldNA. bioRxiv, 2022.2009.2009.507333.

13. Townshend, R.J.L., Eismann, S., Watkins, A.M., Rangan, R., Karelina, M., Das, R. and Dror, R.O. (2021) Geometric deep learning of RNA structure. Science, 373, 1047–1051.

14. Jumper, J., Evans, R., Pritzel, A., Green, T., Figurnov, M., Ronneberger, O., Tunyasuvunakool, K., Bates, R., Zidek, A., Potapenko, A. et al. (2021) Highly accurate protein structure prediction with AlphaFold. Nature, 596, 583–589.

15. Tunyasuvunakool, K., Adler, J., Wu, Z., Green, T., Zielinski, M., Zidek, A., Bridgland, A., Cowie, A., Meyer, C., Laydon, A. et al. (2021) Highly accurate protein structure prediction for the human proteome. Nature, 596, 590–596.

16. van Dijk, M., Thulluru, H.K., Mulders, J., Michel, O.J., Poutsma, A., Windhorst, S., Kleiverda, G., Sie, D., Lachmeijer, A.M.A. and Oudejans, C.B.M. (2012) HELLP babies link a novel lincRNA to the trophoblast cell cycle. J Clin Invest, 122, 4003–4011.

17. Vila-Sanjurjo, A., Ridgeway, W.K., Seymaner, V., Zhang, W., Santoso, S., Yu, K. and Cate, J.H.D. (2003) X-ray crystal structures of the WT and a hyper-accurate ribosome from *Escherichia coli*. Proceedings of the National Academy of Sciences, 100, 8682.

18. Dunkle, J.A., Xiong, L., Mankin, A.S. and Cate, J.H.D. (2010) Structures of the *Escherichia coli* ribosome with antibiotics bound near the peptidyl transferase center explain spectra of drug action. Proceedings of the National Academy of Sciences, 107, 17152.

19. Noeske, J., Wasserman, M.R., Terry, D.S., Altman, R.B., Blanchard, S.C. and Cate, J.H.D. (2015) High-resolution structure of the Escherichia coli ribosome. Nature Structural & Molecular Biology, 22, 336–341.

20. Davis, J.H., Tan, Y.Z., Carragher, B., Potter, C.S., Lyumkis, D. and Williamson, J.R. (2016) Modular Assembly of the Bacterial Large Ribosomal Subunit. Cell, 167, 1610–1622.e1615.

21. Qin, B., Lauer, S.M., Balke, A., Vieira-Vieira, C.H., Bürger, J., Mielke, T., Selbach, M., Scheerer, P., Spahn, C.M.T. and Nikolay, R. (2023) Cryo-EM captures early ribosome assembly in action. Nature Communications, 14, 898.

22. Dai, X., Li, Z., Lai, M., Shu, S., Du, Y., Zhou, Z.H. and Sun, R. (2017) In situ structures of the genome and genome-delivery apparatus in a single-stranded RNA virus. Nature, 541, 112–116.

23. Koning, R.I., Gomez-Blanco, J., Akopjana, I., Vargas, J., Kazaks, A., Tars, K., Carazo, J.M. and Koster, A.J. (2016) Asymmetric cryo-EM reconstruction of phage MS2 reveals genome structure in situ. Nature Communications, 7, 12524.

24. Chang, J.-Y., Gorzelnik, K.V., Thongchol, J. and Zhang, J. (2022) Structural Assembly of Qβ Virion and Its Diverse Forms of Virus-like Particles. Viruses, 10.3390/v14020225.

25. Ma, H., Pham, P., Luo, B., Rangan, R., Kappel, K., Su, Z. and Das, R. (2023) In Ding, J., Stagno, J. R. and Wang, Y.-X. (eds.), *RNA Structure and Dynamics*. Springer US, New York, NY, pp. 193–211.

26. Kappel, K., Liu, S., Larsen, K.P., Skiniotis, G., Puglisi, E.V., Puglisi, J.D., Zhou, Z.H., Zhao, R. and Das, R. (2018) De novo computational RNA modeling into cryo-EM maps of large ribonucleoprotein complexes. Nature Methods, 15, 947–954.

27. Fang, X., Stango, J.R., Bhandari, Y.R., Zuo, X., Wang, Y.,. (2015) Small-angle X-ray scattering: a bridge between RNA secondary structures and three-dimensional topological structures. Structural Biology, 30, 147–160.

28. Gopal, A., Zhou, Z.H., Knobler, C.M. and Gelbart, W.M. (2012) Visualizing large RNA molecules in solution. RNA (New York, N.Y.), 18, 284–299.

29. San Emeterio, J. and Pollack, L. (2020) Visualizing a viral genome with contrast variation small angle X-ray scattering. Journal of Biological Chemistry, 295, 15923–15932.

30. Garmann, R.F., Gopal, A., Athavale, S.S., Knobler, C.M., Gelbart, W.M. and Harvey, S.C. (2015) Visualizing the global secondary structure of a viral RNA genome with cryo-electron microscopy. RNA (New York, N.Y.), 21, 877–886.

31. Zeng, Y., Larson, S.B., Heitsch, C.E., McPherson, A. and Harvey, S.C. (2012) A model for the structure of satellite tobacco mosaic virus. Journal of structural biology, 180, 110–116.

32. Poblete, S. and Guzman, H.V. (2021) Structural 3D Domain Reconstruction of the RNA Genome from Viruses with Secondary Structure Models. Viruses, 13.

33. Popenda, M., Szachniuk, M., Antczak, M., Purzycka, K.J., Lukasiak, P., Bartol, N., Blazewicz, J. and Adamiak, R.W. (2012) Automated 3D structure composition for large RNAs. Nucleic acids research, 40, e112–e112.

34. Jonikas, M.A., Radmer, R.J., Laederach, A., Das, R., Pearlman, S., Herschlag, D. and Altman, R.B. (2009) Coarse-grained modeling of large RNA molecules with knowledge-based potentials and structural filters. RNA (New York, N.Y.), 15, 189–199.

35. Popenda, M., Zok, T., Sarzynska, J., Korpeta, A., Adamiak, Ryszard W., Antczak, M. and Szachniuk, M. (2021) Entanglements of structure elements revealed in RNA 3D models. Nucleic Acids Research, 49, 9625–9632.

36. Elcock, A.H. (2023) Easy Removal of Steric Clashes and Entanglements in Macromolecular Systems by Temporary Addition of a Fourth Spatial Dimension (preprint). bioRxiv.

37. Frembgen-Kesner, T. and Elcock, A.H. (2009) Striking Effects of Hydrodynamic Interactions on the Simulated Diffusion and Folding of Proteins. Journal of Chemical Theory and Computation, 5, 242–256.

38. Benítez, A.A., Hernández Cifre, J.G., Díaz Baños, F.G. and de la Torre, J.G. (2015) Prediction of solution properties and dynamics of RNAs by means of Brownian dynamics simulation of coarse-grained models: Ribosomal 5S RNA and phenylalanine transfer RNA. BMC Biophysics, 8, 11.

39. Dawson, W.K., Maciejczyk, M., Jankowska, E.J. and Bujnicki, J.M. (2016) Coarse-grained modeling of RNA 3D structure. Methods, 103, 138–156.

40. Li, J. and Chen, S.-J. (2021) RNA 3D Structure Prediction Using Coarse-Grained Models. Frontiers in Molecular Biosciences, 8.

41. Hacker, W.C., Li, S. and Elcock, A.H. (2017) Features of genomic organization in a nucleotide-resolution molecular model of the Escherichia coli chromosome. Nucleic Acids Research, 45, 7541–7554.

42. Malhotra, A., Tan, R.K. and Harvey, S.C. (1994) Modeling large RNAs and ribonucleoprotein particles using molecular mechanics techniques. Biophysical Journal, 66, 1777–1795.

43. Hyeon, C., Dima, R.I. and Thirumalai, D. (2006) Pathways and Kinetic Barriers in Mechanical Unfolding and Refolding of RNA and Proteins. Structure, 14, 1633–1645.

44. Helmling, C., Keyhani, S., Sochor, F., Fürtig, B., Hengesbach, M. and Schwalbe, H. (2015) Rapid NMR screening of RNA secondary structure and binding. Journal of Biomolecular NMR, 63, 67–76.

45. Madden Emily, A., Plante Kenneth, S., Morrison Clayton, R., Kutchko Katrina, M., Sanders, W., Long Kristin, M., Taft-Benz, S., Cruz Cisneros Marta, C., White Ashlyn, M., Sarkar, S., et al. Using SHAPE-MaP To Model RNA Secondary Structure and Identify 3’UTR Variation in Chikungunya Virus. Journal of Virology, 94, e00701–00720.

46. Larman, B.C., Dethoff, E.A. and Weeks, K.M. (2017) Packaged and Free Satellite Tobacco Mosaic Virus (STMV) RNA Genomes Adopt Distinct Conformational States. Biochemistry, 56, 2175–2183.

47. Zubradt, M., Gupta, P., Persad, S., Lambowitz, A.M., Weissman, J.S. and Rouskin, S. (2017) DMS-MaPseq for genome-wide or targeted RNA structure probing in vivo. Nature Methods, 14, 75–82.

48. Hofacker, I.L. (2003) Vienna RNA secondary structure server. Nucleic acids research, 31, 3429–3431.

49. Proctor, J.R. and Meyer, I.M. (2013) COFOLD: an RNA secondary structure prediction method that takes co-transcriptional folding into account. Nucleic acids research, 41, e102–e102.

50. Mathews, D.H., Disney, M.D., Childs, J.L., Schroeder, S.J., Zuker, M. and Turner, D.H. (2004) Incorporating chemical modification constraints into a dynamic programming algorithm for prediction of RNA secondary structure. Proceedings of the National Academy of Sciences of the United States of America, 101, 7287.

51. Turner, D.H. and Mathews, D.H. (2010) NNDB: the nearest neighbor parameter database for predicting stability of nucleic acid secondary structure. Nucleic acids research, 38, D280–D282.

52. Lorenz, R., Bernhart, S.H., Höner zu Siederdissen, C., Tafer, H., Flamm, C., Stadler, P.F. and Hofacker, I.L. (2011) ViennaRNA Package 2.0. Algorithms for Molecular Biology, 6, 26.

53. Bernhart, S.H., Tafer, H., Mückstein, U., Flamm, C., Stadler, P.F. and Hofacker, I.L. (2006) Partition function and base pairing probabilities of RNA heterodimers. Algorithms for Molecular Biology, 1, 3.

54. Hofacker, I.L., Fontana, W., Stadler, P.F., Bonhoeffer, L.S., Tacker, M. and Schuster, P. (1994) Fast folding and comparison of RNA secondary structures. Monatshefte für Chemie / Chemical Monthly, 125, 167–188.

55. Lorenz, R., Hofacker, I.L. and Stadler, P.F. (2016) RNA folding with hard and soft constraints. Algorithms for Molecular Biology, 11, 8.

56. Tworek, J.W. and Elcock, A.H. (2023) An orientationally averaged version of the Rotne-Prager-Yamakawa tensor provides a fast but still accurate treatment of hydrodynamic interactions in Brownian dynamics simulations of biological macromolecules (preprint). bioRxiv.

57. Cheatham, T.E., 3rd, Brooks, B.R. and Kollman, P.A. (2001) Molecular modeling of nucleic acid structure. Curr Protoc Nucleic Acid Chem, Chapter 7, Unit-7.5.

58. Kondo, J., Tada, Y., Dairaku, T., Saneyoshi, H., Okamoto, I., Tanaka, Y. and Ono, A. (2015) High-Resolution Crystal Structure of a Silver(I)–RNA Hybrid Duplex Containing Watson–Crick-like C Silver(I) Metallo-Base Pairs. Angewandte Chemie International Edition, 54, 13323–13326.

59. Holbrook, S.R., Cheong, C., Tinoco, I. and Kim, S.-H. (1991) Crystal structure of an RNA double helix incorporating a track of non-Watson–Crick base pairs. Nature, 353, 579–581.

60. Möller, T. and Trumbore, B. (1997) Fast, Minimum Storage Ray-Triangle Intersection. Journal of Graphics Tools, 2, 21–28.

61. Yamakawa, H. (1970) Transport Properties of Polymer Chains in Dilute Solution: Hydrodynamic Interaction. The Journal of Chemical Physics, 53, 436–443.

62. Rotne, J. and Prager, S. (1969) Variational Treatment of Hydrodynamic Interaction in Polymers. The Journal of Chemical Physics, 50, 4831–4837.

63. Abraham, M.J., Murtola, T., Schulz, R., Páll, S., Smith, J.C., Hess, B. and Lindahl, E. (2015) GROMACS: High performance molecular simulations through multi-level parallelism from laptops to supercomputers. SoftwareX, 1, 19–25.

64. Werner, A. (2011) Predicting translational diffusion of evolutionary conserved RNA structures by the nucleotide number. Nucleic acids research, 39, e17–e17.

65. Berman, H.M., Westbrook, J., Feng, Z., Gilliland, G., Bhat, T.N., Weissig, H., Shindyalov, I.N. and Bourne, P.E. (2000) The Protein Data Bank. Nucleic acids research, 28, 235–242.

66. Kapral, G.J., Jain, S., Noeske, J., Doudna, J.A., Richardson, D.C. and Richardson, J.S. (2014) New tools provide a second look at HDV ribozyme structure, dynamics and cleavage. Nucleic Acids Research, 42, 12833–12846.

67. B. Rupp, S.P. (1996) PDBSUP—a FORTRAN program that determines the rotation matrix and translation vector for best fit superposition of two pdb files by solving the quaternion eigenvalue problem. Lawrence Livermore National Laboratory, Livermore, CA.

68. Schultes, E.A., Spasic, A., Mohanty, U. and Bartel, D.P. (2005) Compact and ordered collapse of randomly generated RNA sequences. Nature Structural & Molecular Biology, 12, 1130–1136.

69. Borodavka, A., Singaram, S.W., Stockley, P.G., Gelbart, W.M., Ben-Shaul, A. and Tuma, R. (2016) Sizes of Long RNA Molecules Are Determined by the Branching Patterns of Their Secondary Structures. Biophysical journal, 111, 2077–2085.

70. Antczak, M., Zok, T., Popenda, M., Lukasiak, P., Adamiak, R.W., Blazewicz, J. and Szachniuk, M. (2014) RNApdbee--a webserver to derive secondary structures from pdb files of knotted and unknotted RNAs. Nucleic acids research, 42, W368–W372.

71. Zok, T., Antczak, M., Zurkowski, M., Popenda, M., Blazewicz, J., Adamiak, R.W. and Szachniuk, M. (2018) RNApdbee 2.0: multifunctional tool for RNA structure annotation. Nucleic Acids Research, 46, W30–W35.

72. Loveland, A.B., Demo, G., Grigorieff, N. and Korostelev, A.A. (2017) Ensemble cryo-EM elucidates the mechanism of translation fidelity. Nature, 546, 113–117.

73. Burley, S.K., Berman, H.M., Christie, C., Duarte, J.M., Feng, Z., Westbrook, J., Young, J. and Zardecki, C. (2018) RCSB Protein Data Bank: Sustaining a living digital data resource that enables breakthroughs in scientific research and biomedical education. Protein science : a publication of the Protein Society, 27, 316–330.

74. Bhattacharjee, S.M., Giacometti, A. and Maritan, A. (2013) Flory theory for polymers. Journal of Physics: Condensed Matter, 25, 503101.

75. Yoffe, A.M., Prinsen, P., Gelbart, W.M. and Ben-Shaul, A. (2011) The ends of a large RNA molecule are necessarily close. Nucleic acids research, 39, 292–299.

76. Jonikas, M.A., Radmer, R.J. and Altman, R.B. (2009) Knowledge-based instantiation of full atomic detail into coarse-grain RNA 3D structural models. Bioinformatics (Oxford, England), 25, 3259–3266.

77. Ando, T., Chow, E., Saad, Y. and Skolnick, J. (2012) Krylov subspace methods for computing hydrodynamic interactions in Brownian dynamics simulations. The Journal of chemical physics, 137, 064106.

78. Fernandes, M.X., Ortega, A., López Martínez, M.C. and García de la Torre, J. (2002) Calculation of hydrodynamic properties of small nucleic acids from their atomic structure. Nucleic Acids Research, 30, 1782–1788.

79. Folkhard, W., Pilz, I., Kratky, O., Garrett, R. and StÖFfler, G. (1975) Small-Angle X-Ray Studies on the Structure of 16-S Ribosomal RNA and of a Complex of Ribosomal Protein S4 and 16-S Ribosomal RNA from Escherichia coli. Eur J Biochem, 59, 63–71.

80. Timchenko, A.A., Langowski, J. and Serdyuk, I.N. (1993) Structural changes in 16S RNA from Escherichia coli upon unfolding by urea. Biopolymers, 33, 1747–1755.

81. Tam, M.F., Dodd, J.A. and Hill, W.E. (1981) Physical characteristics of 16 S rRNA under reconstitution conditions. Journal of Biological Chemistry, 256, 6430–6434.

82. Hill, W.E., Bakke, K.R. and Blair, D.P. (1977) On the sedimentation behavior and molecular weight of 16S ribosomal RNA from Escherichia coli. Nucleic Acids Research, 4, 473–476.

83. Allen, S.H. and Wong, K.-P. (1986) The role of magnesium and potassium ions in the molecular mechanism of ribosome assembly: Hydrodynamic, conformational, and thermal stability studies of 16 S RNA from Escherichia coli ribosomes. Archives of Biochemistry and Biophysics, 249, 137–147.

84. Singh, J., Paliwal, K., Zhang, T., Singh, J., Litfin, T. and Zhou, Y. (2021) Improved RNA secondary structure and tertiary base-pairing prediction using evolutionary profile, mutational coupling and two-dimensional transfer learning. Bioinformatics, 37, 2589–2600.

85. Singh, J., Hanson, J., Paliwal, K. and Zhou, Y. (2019) RNA secondary structure prediction using an ensemble of two-dimensional deep neural networks and transfer learning. Nature Communications, 10, 5407.

86. Humphrey, W., Dalke, A. and Schulten, K. (1996) VMD: visual molecular dynamics. Journal of molecular graphics, 14, 33–38.

