## Supporting Information for "Modeling the 3D structure and conformational dynamics of very large RNAs using coarse-grained molecular simulations"

**for:**

### Supporting Table & Figure Legends

**Table S1** **Simulated hydrodynamic radii for all 21 85-nt RNA.** All Rh values are reported in Ångstroms. Error bars represent the standard deviation of Rh values calculated from three contiguous blocks of the 10 µs simulation (i.e. 1.00-3.33 µs, 3.34-6.66 µs, 6.67-10.0 µs).

**Table S2** **Computed shape anisotropies for 16 large RNAs.** All Rh values are reported in Ångstroms. Error bars represent the standard deviation of Rh values calculated from three contiguous blocks of the 10 µs simulation (i.e. 1.00-3.33 µs, 3.34-6.66 µs, 6.67-10.0 µs).

**Table S3** **Computed shape anisotropies for 16 large RNAs.** All Rh values are reported in Ångstroms. Error bars represent the standard deviation of Rh values calculated from three contiguous blocks of the 10 µs simulation (i.e. 1.00-3.33 µs, 3.34-6.66 µs, 6.67-10.0 µs).

**Figure S1** **Resolution of entanglements by temporary addition of a 4$^{th}$ spatial dimension.** Images showing how an entanglement of two helical regions in a single RNA structure is automatically resolved by the modelling protocol presented here. The top panel shows how two stem-loops have been inadvertently entangled during the first two stages of the protocol. Prior to the third stage, all elements of secondary structure are assigned a different displacement in a temporary 4$^{th}$ spatial dimension such that all of them are suitably separated from each other when their separation distances are calculated in all 4 dimensions. As the simulation proceeds, and the 4$^{th}$ spatial dimension is reduced in extent, secondary structure elements are forced to approach and accommodate each other in the remaining 3 dimensions. In the case shown, the two stem-loops automatically move apart from each other.

**Figure S2** **BD-HI simulations of structured RNAs non-trivially reproduce experimental hydrodynamic radii. A.** Scatter plot comparing hydrodynamic radii, Rh, computed from BD-HI simulations of five structured RNAs with their corresponding experimental values; the line represents the linear regression ($r^2$ = 0.91). **B.** Scatter plot comparing the experimental hydrodynamic radii, Rh, of the same RNAs with the number of nucleotides; the line represents the regression of the data to a power law ($r^2$ = 0.57). **C.** Scatter plot comparing hydrodynamic radii, Rh, computed from BD-HI simulations of the same RNAs in their unfolded states with the experimental folded state values; the line represents the linear regression ($r^2$ = 0.57). **D.** Scatter plot comparing hydrodynamic radii, Rh, computed from BD-HI simulations of the same RNAs in their unfolded states with the number of nucleotides; the line represents the regression of the data to a power law ($r^2$ = 0.98).

**Figure S3** **Application of the 3D modeling protocol with RNAFold to large RNAs.** Images of all 16 3D models built using RNAFold-predicted secondary structures at the end of a 10 µs BD-HI simulation.

**Figure S4** **Application of the 3D modeling protocol with CoFold to large RNAs.** Images of all 16 3D models built using CoFold-predicted secondary structures at the end of a 10 µs BD-HI simulation.

**Figure S5** **Simulated hydrodynamic radii of large RNAs scale with the number of nucleotides. A.** Scatter plot comparing hydrodynamic radii, Rh, computed from BD-HI simulations of 16 large RNAs using RNAFold-predicted secondary structures with those obtained using CoFold-predicted secondary structures; the line represents y = x. **B.** Scatter plot comparing hydrodynamic radii, Rh, computed from BD-HI simulations of the 16 large RNAs (built using RNAFold-predicted secondary structures) with the number of nucleotides; the line represents the regression of the data to a power law ($r^2$ = 0.87). **C.** Scatter plot comparing hydrodynamic radii, Rh, computed from BD-HI simulations of the 16 large RNAs (built using CoFold-predicted secondary structures) with the number of nucleotides; the line represents the regression of the data to a power law ($r^2$ = 0.94).

**Figure S6** **Estimates of the 3D models' hydrodynamic radii converge within 10 µs BD-HI simulation.** Plots of the hydrodynamic radii, Rh, computed from BD-HI simulations from three contiguous blocks of the 10 µs simulation (i.e. 1.00-3.33 µs, 3.34-6.66 µs, 6.67-10.0 µs). **A.** Rh values for each block for 3D models built using RNAFold-predicted secondary structures. **B.** Same as **A** but for 3D models built using CoFold-predicted secondary structures.

**Figure S7** **Simulated hydrodynamic radii are insensitive to modest variations in the secondary structure predictions.** A. Images of 5 alternative 3D models of the 961-nt BunVS RNA (built using different RNAFold-predicted secondary structures) at the end of a 10 µs BD-HI simulation. **B.** Same as **A** but showing alternative 3D models of the 1400-nt FHV2 RNA. **C.** Bar chart illustrating the hydrodynamic radii, Rh, computed from BD-HI simulations of the 5 alternative 3D models and comparing them with the values obtained from the corresponding minimum free-energy secondary structure prediction.

**Figure S8** **Structure of the predicted 16S rRNA dimer predicted by RNAcofold.** The two copies of the 16S rRNA are shown in red and blue.

**Figure S9** **Shape anisotropy calculations show behavior mirroring data from cryo-EM data.** Scatter plots of the maximum diameter (M) and minimum diameter (m) versus the anisotropy, A, calculated for snapshots sampled from 10 µs BD-HI simulations. **A.** Results from the Gelbart group's 975-nt RNA obtained from a 3D model built using an RNAFold-predicted secondary structure. **B.** Same as **A** but for the Gelbart group's 1523-nt RNA. **C.** Same as **A** but for a 3D model built using a CoFold-predicted secondary structure. **D.** Same as **C** but for the Gelbart group's 1523-nt RNA.

**Figure S10** **Estimates of the 3D models' shape anisotropy converge within 10 µs BD-HI simulation.** Plots of the cumulative mean anisotropy, A, computed from 10 µs BD-HI simulations as a function of increasing simulation time. **A.** Mean anisotropy of the Gelbart group's 975-nt RNA obtained from a 3D model built using an RNAFold-predicted secondary structure. **B.** Same as **A** but for the Gelbart group's 1523-nt RNA. **C.** Same as **A** but for a 3D model built using a CoFold-predicted secondary structure. **D.** Same as **C** but for the Gelbart group's 1523-nt RNA.

**Figure S11** **Comparison with experiment is excellent for large RNAs that show $Mg^{2+}$-dependent compaction.** Same as Figure 4C of the main text but showing results only for RNAs show Rh values show a significant dependence on $Mg^{2+}$ in the experiments of Borodavka et al.

| RNA # | I series | P series | $U_{85}$ |
|---|---|---|---|
| 1 | 31.5 ± 1.4 | 29.8 ± 1.9 | 41.0 ± 2.4 |
| 2 | 33.0 ± 1.9 | 32.5 ± 1.8 | |
| 3 | 32.1 ± 2.7 | 30.8 ± 1.6 | |
| 4 | 31.7 ± 1.4 | 31.9 ± 1.4 | |
| 5 | 33.5 ± 2.0 | 31.8 ± 2.1 | |
| 6 | 32.7 ± 1.1 | 32.1 ± 1.0 | |
| 7 | 32.6 ± 1.8 | 30.7 ± 1.2 | |
| 8 | 31.3 ± 1.4 | 32.7 ± 2.1 | |
| 9 | 31.3 ± 2.0 | 29.3 ± 1.5 | |
| 10 | 31.3 ± 1.3 | 30.7 ± 1.4 | |

**Table S1**

| **RNA name** | **<A>** | **a:b:c** |
|---|---|---|
| RV s11 | 0.34 | 5.4:3.2:1 |
| RV s11 scrambled | 0.28 | 4.3:3.1:1 |
| BunVS | 0.24 | 2.9:2.9:1 |
| STNV | 0.22 | 3.9:3.8:1 |
| FHV 2 | 0.27 | 3.9:2.9:1 |
| 16S rRNA | 0.22 | 2.9:2.4:1 |
| Ef1-alpha mRNA | 0.25 | 3.2:2.4:1 |
| HOX lncRNA | 0.27 | 3.7:2.7:1 |
| 5’-MS2 | 0.28 | 3.4:2.2:1 |
| 3’-MS2 | 0.14 | 2.0:2.0:1 |
| NRON | 0.21 | 3.0:2.8:1 |
| 23S rRNA | 0.20 | 3.0:2.8:1 |
| FHV 1 | 0.21 | 2.6:2.1:1 |
| RV s1 | 0.25 | 3.8:3.1:1 |
| MS2 | 0.20 | 2.8:2.5:1 |
| RpoB mRNA | 0.17 | 3.2:3.2:1 |
| **AVERAGE** | **0.23** | **3.4:2.8:1** |

**Table S2**

| **RNA name** | **<A>** | **a:b:c** |
|---|---|---|
| RV s11 | 0.22 | 4.1:4.0:1 |
| RV s11 scrambled | 0.24 | 4.4:4.0:1 |
| BunVS | 0.26 | 3.8:3.0:1 |
| STNV | 0.27 | 4.5:3.5:1 |
| FHV 2 | 0.23 | 4.2:3.9:1 |
| 16S rRNA | 0.24 | 3.3:2.6:1 |
| Ef1-alpha mRNA | 0.21 | 3.1:2.9:1 |
| HOX lncRNA | 0.23 | 3.4:2.2:1 |
| 5’-MS2 | 0.35 | 4.5:2.4:1 |
| 3’-MS2 | 0.29 | 4.0:2.7:1 |
| NRON | 0.33 | 4.1:2.3:1 |
| 23S rRNA | 0.28 | 4.0:2.8:1 |
| FHV 1 | 0.29 | 4.6:3.3:1 |
| RV s1 | 0.34 | 4.7:2.7:1 |
| MS2 | 0.26 | 3.7:2.8:1 |
| RpoB mRNA | 0.33 | 3.9:3.3:1 |
| **AVERAGE** | **0.27** | **4.0:3.0:1** |

### Table S3

**Figure S1**

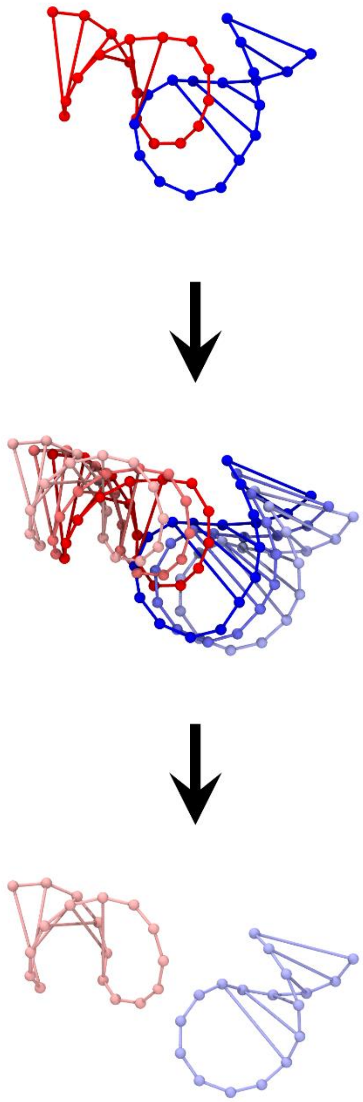

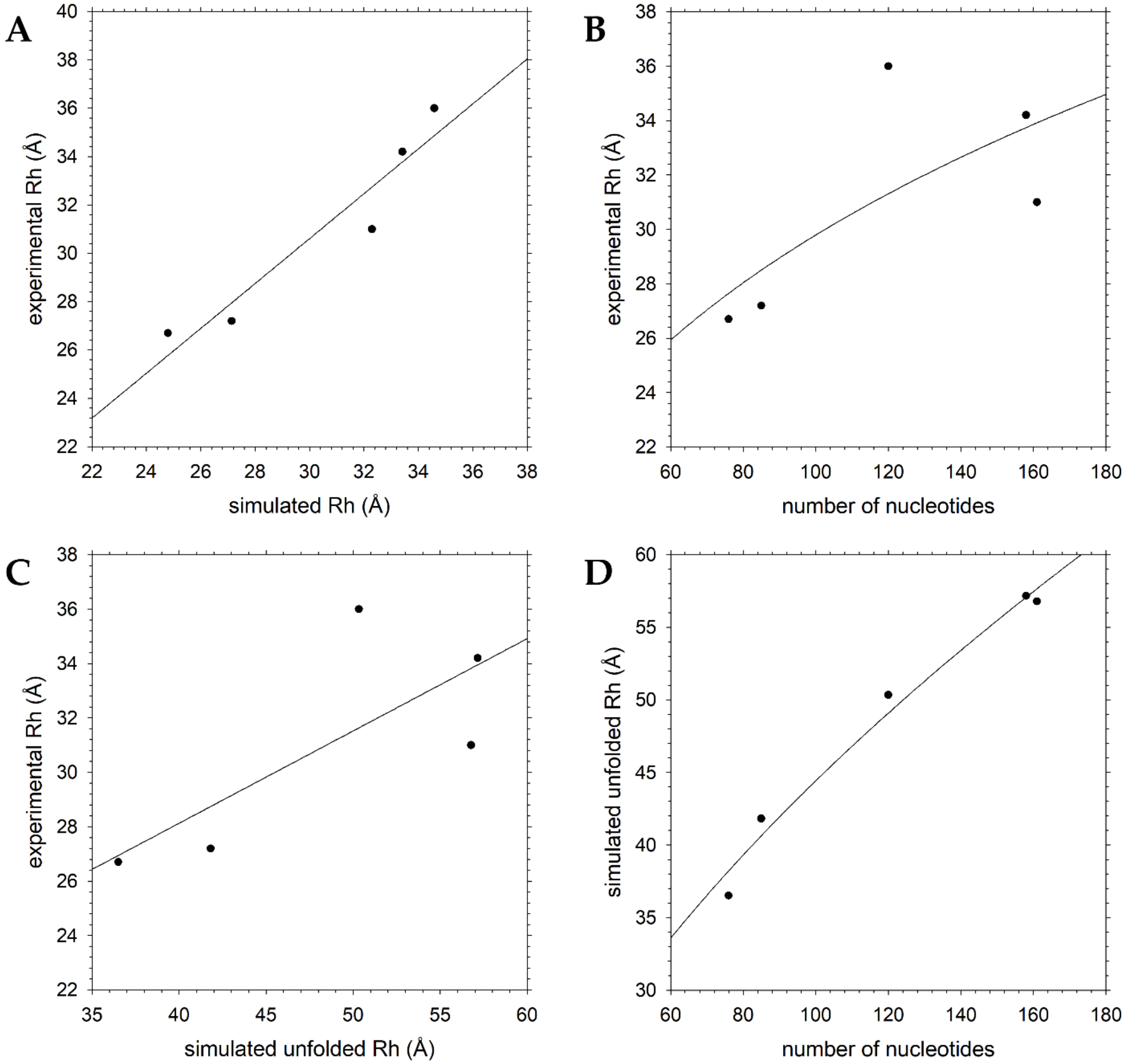
A
B
C
D
experimental Rh (Å)
simulated Rh (Å)
number of nucleotides
simulated unfolded Rh (Å)


**Figure S2**

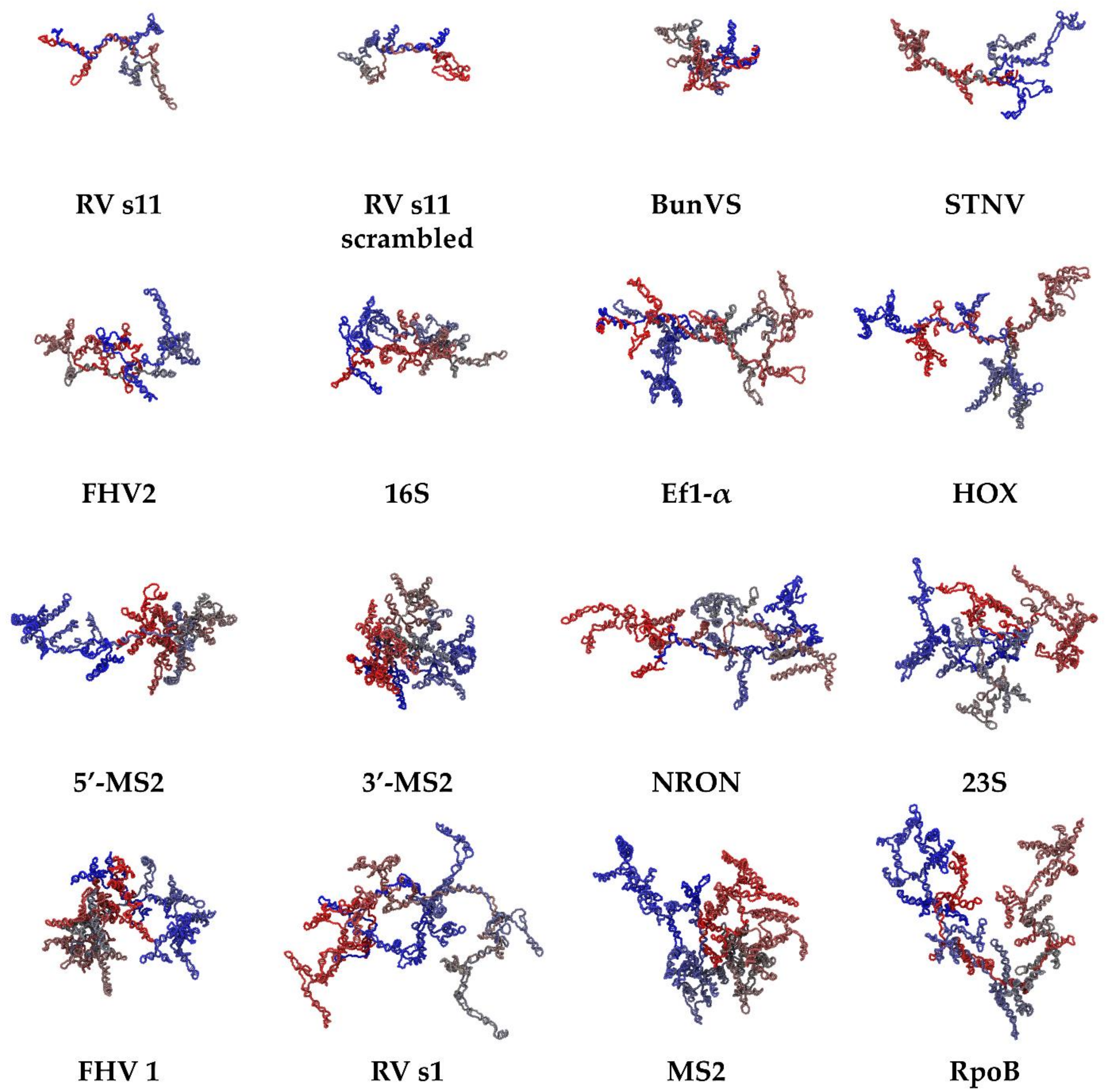

RV s11
RV s11 scrambled
BunVS
STNV
FHV2
16S
Ef1-$\alpha$
HOX
5′-MS2
3′-MS2
NRON
23S
FHV 1
RV s1
MS2
RpoB


**Figure S3**

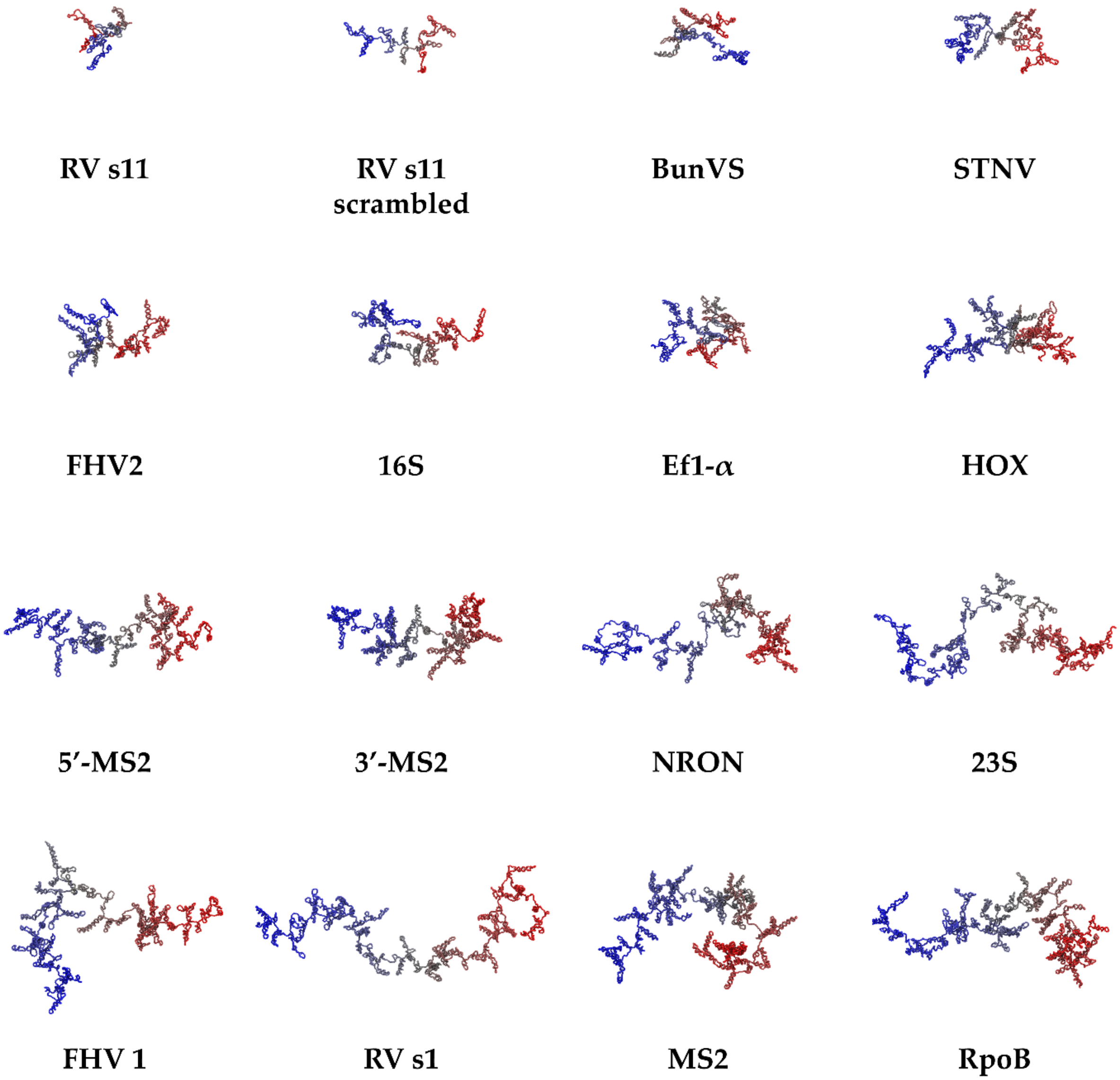
RV s11
RV s11 scrambled
BunVS
STNV
FHV2
16S
Ef1-α
HOX
5′-MS2
3′-MS2
NRON
23S
FHV 1
RV s1
MS2
RpoB

**Figure S4**

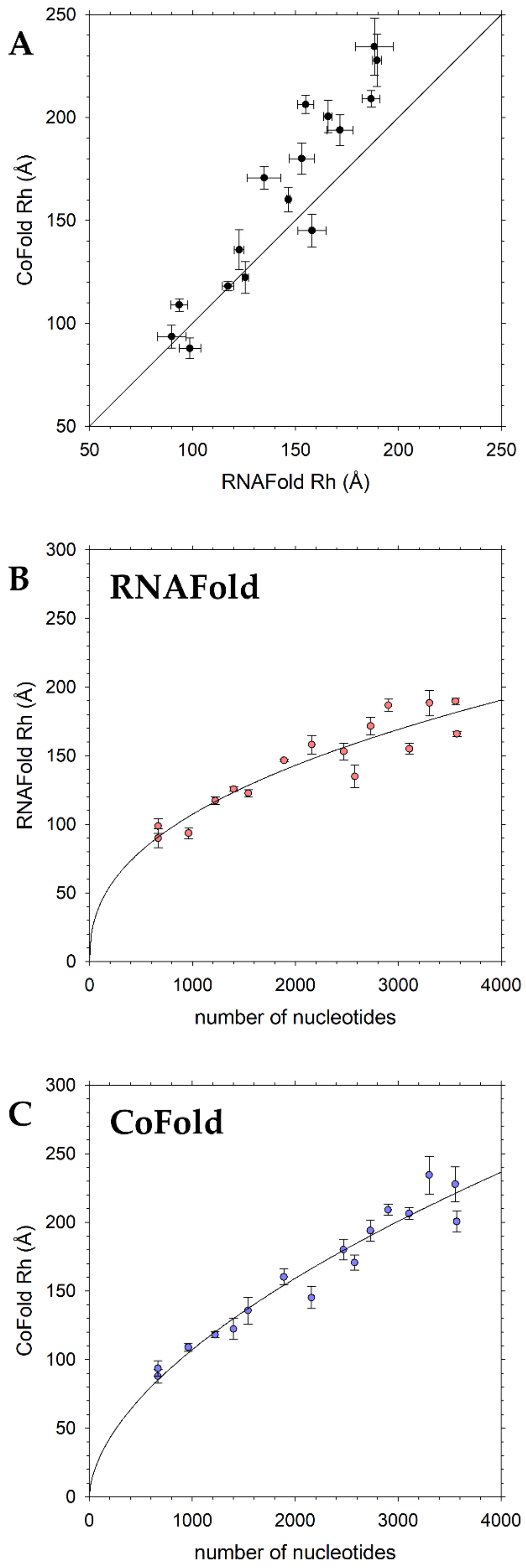
A
CoFold Rh (Å)
RNAFold Rh (Å)
250
200
150
100
50
50
100
150
200
250
B
RNAFold
RNAFold Rh (Å)
number of nucleotides
300
250
200
150
100
50
0
0
1000
2000
3000
4000
C
CoFold
CoFold Rh (Å)
number of nucleotides
300
250
200
150
100
50
0
0
1000
2000
3000
4000

**Figure S5**

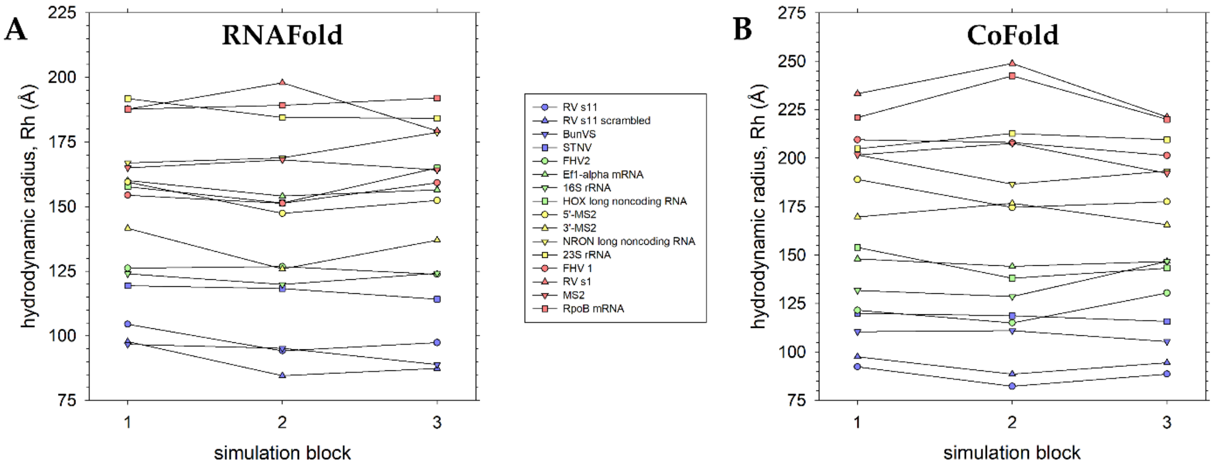
A
RNAFold
B
CoFold
hydrodynamic radius, Rh (Å)
simulation block
RV s11
RV s11 scrambled
BunVS
STNV
FHV2
Ef1-alpha mRNA
16S rRNA
HOX long noncoding RNA
5'-MS2
3'-MS2
NRON long noncoding RNA
23S rRNA
FHV 1
RV s1
MS2
RpoB mRNA

**Figure S6**

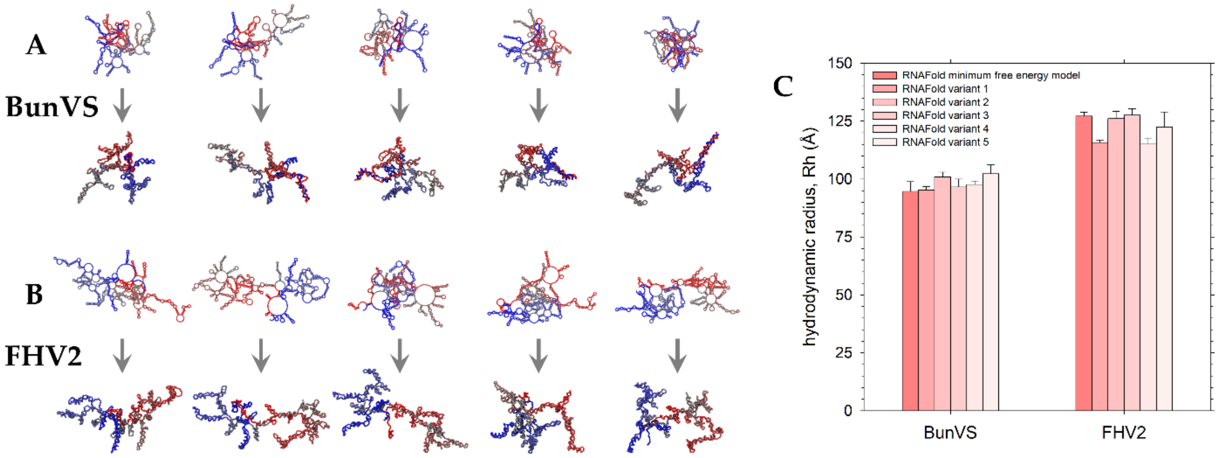
A
BunVS
B
FHV2
C
RNAFold minimum free energy model
RNAFold variant 1
RNAFold variant 2
RNAFold variant 3
RNAFold variant 4
RNAFold variant 5
hydrodynamic radius, Rh (Å)
150
125
100
75
50
25
0
BunVS
FHV2

**Figure S7**

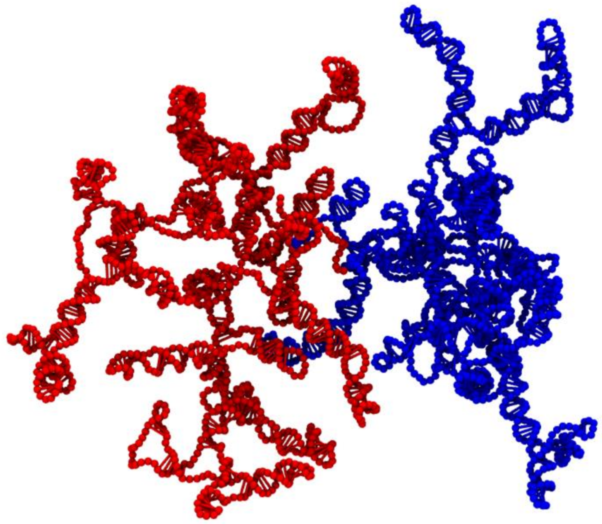

**Figure S8**

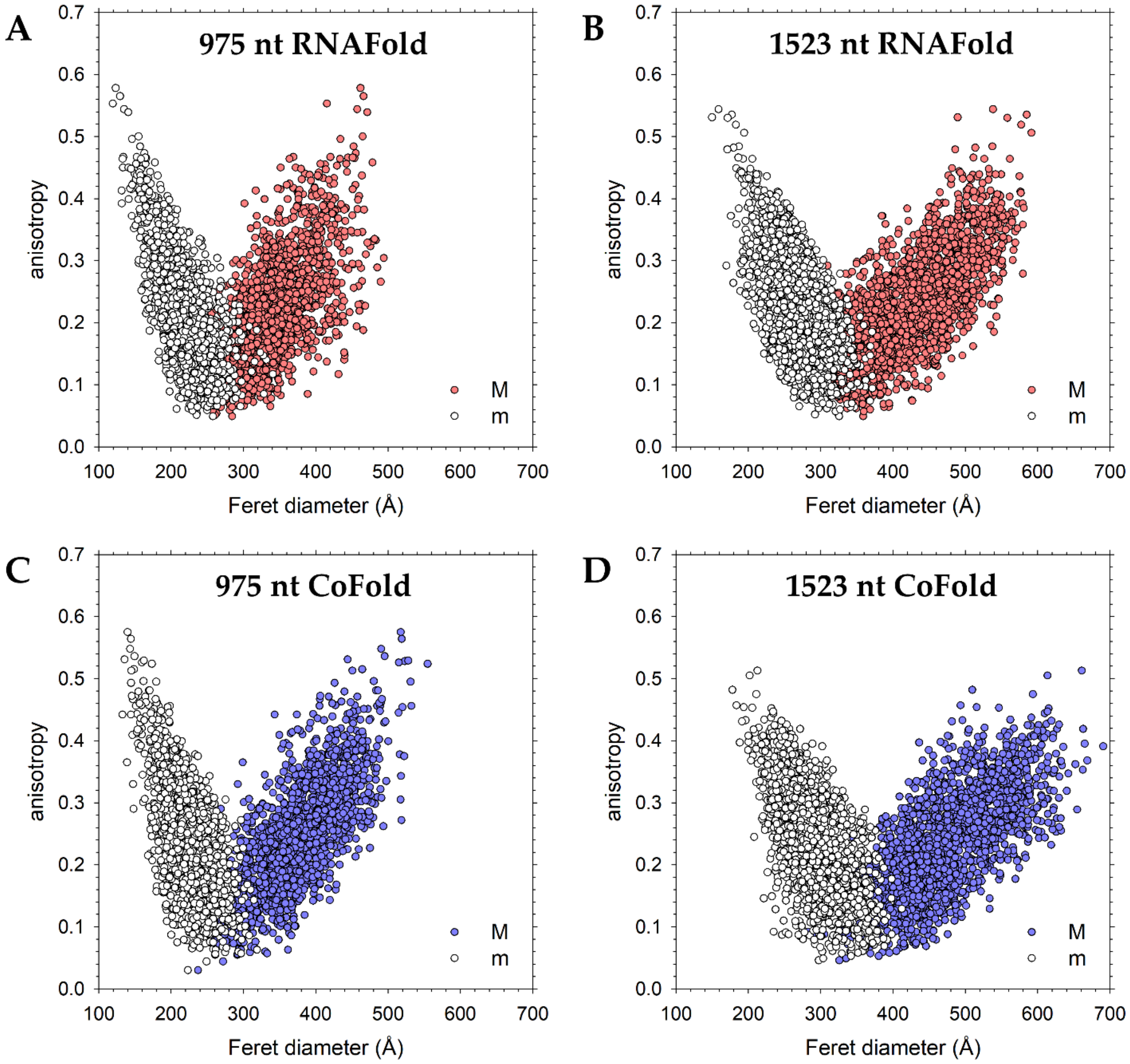
A
975 nt RNAFold
anisotropy
Feret diameter (Å)
M
m
B
1523 nt RNAFold
C
975 nt CoFold
D
1523 nt CoFold

**Figure S9**

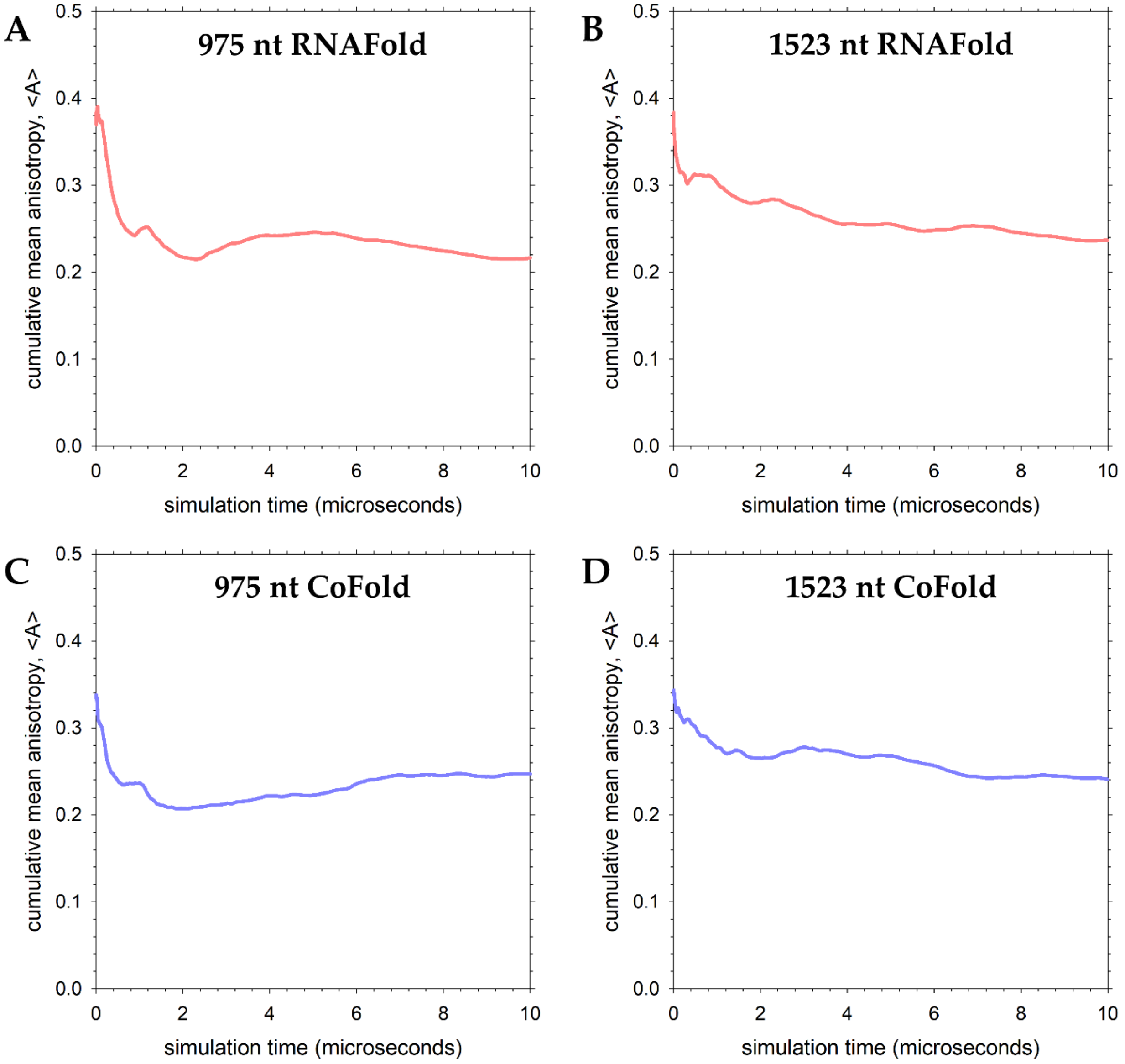
A
975 nt RNAFold
B
1523 nt RNAFold
C
975 nt CoFold
D
1523 nt CoFold
cumulative mean anisotropy, <A>
simulation time (microseconds)
0.0
0.1
0.2
0.3
0.4
0.5
0
2
4
6
8
10

**Figure S10**

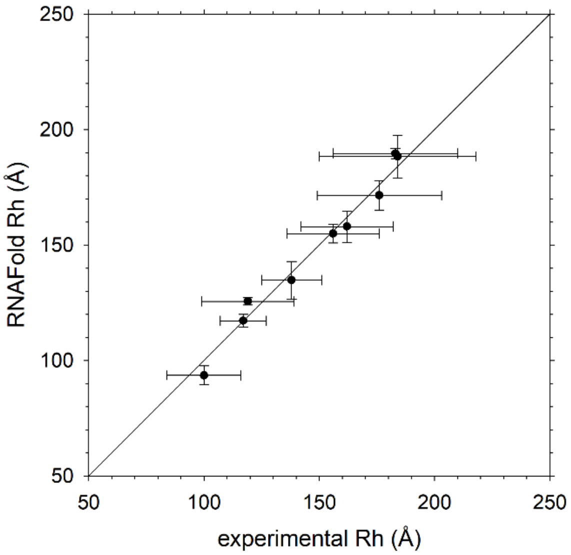
250
200
150
100
50
50
100
150
200
250
RNAFold Rh (Å)
experimental Rh (Å)

**Figure S11**
